# Flexibility and distributive synthesis regulate RNA priming and handoff in human DNA polymerase α-primase

**DOI:** 10.1101/2023.08.01.551538

**Authors:** John J. Cordoba, Elwood A. Mullins, Lauren E. Salay, Brandt F. Eichman, Walter J. Chazin

## Abstract

DNA replication in eukaryotes relies on the synthesis of a ∼30-nucleotide RNA/DNA primer strand through the dual action of the heterotetrameric polymerase α-primase (pol-prim) enzyme. Synthesis of the 7-10-nucleotide RNA primer is regulated by the C-terminal domain of the primase regulatory subunit (PRIM2C) and is followed by intramolecular handoff of the primer to pol α for extension by ∼20 nucleotides of DNA. Here we provide evidence that RNA primer synthesis is governed by a combination of the high affinity and flexible linkage of the PRIM2C domain and the low affinity of the primase catalytic domain (PRIM1) for substrate. Using a combination of small angle X-ray scattering and electron microscopy, we found significant variability in the organization of PRIM2C and PRIM1 in the absence and presence of substrate, and that the population of structures with both PRIM2C and PRIM1 in a configuration aligned for synthesis is low. Crosslinking was used to visualize the orientation of PRIM2C and PRIM1 when engaged by substrate as observed by electron microscopy. Microscale thermophoresis was used to measure substrate affinities for a series of pol-prim constructs, which showed that the PRIM1 catalytic domain does not bind the template or emergent RNA-primed templates with appreciable affinity. Together, these findings support a model of RNA primer synthesis in which generation of the nascent RNA strand and handoff of the RNA-primed template from primase to polymerase α is mediated by the high degree of inter-domain flexibility of pol-prim, the ready dissociation of PRIM1 from its substrate, and the much higher affinity of the POLA1cat domain of polymerase α for full-length RNA-primed templates.

## Introduction

In eukaryotes, DNA polymerase α-primase (pol-prim) initiates DNA synthesis during replication, generating the first ∼30 nucleotides of nascent strands.^1–5^ Pol-prim is unique among replicative polymerases in its ability to perform de novo synthesis from a single-stranded DNA template; the primers it creates are required for further synthesis by the processive polymerases ε and δ that perform the bulk of nascent strand synthesis^6,7^. Pol-prim plays a particularly crucial role on the lagging strand during replication, as primers must be repeatedly synthesized due to the discontinuous nature of synthesis of this nascent strand.^8^ The primers generated by pol-prim are chimeric in nature, consisting of 7-10 ribonucleotides followed by ∼20 deoxyribonucleotides.^9,10^ The RNA and DNA portions of this chimeric primer are generated by distinct active sites located in the primase and polymerase subunits of this tetrameric enzyme (Figure 1).^11^

**Figure 1.**
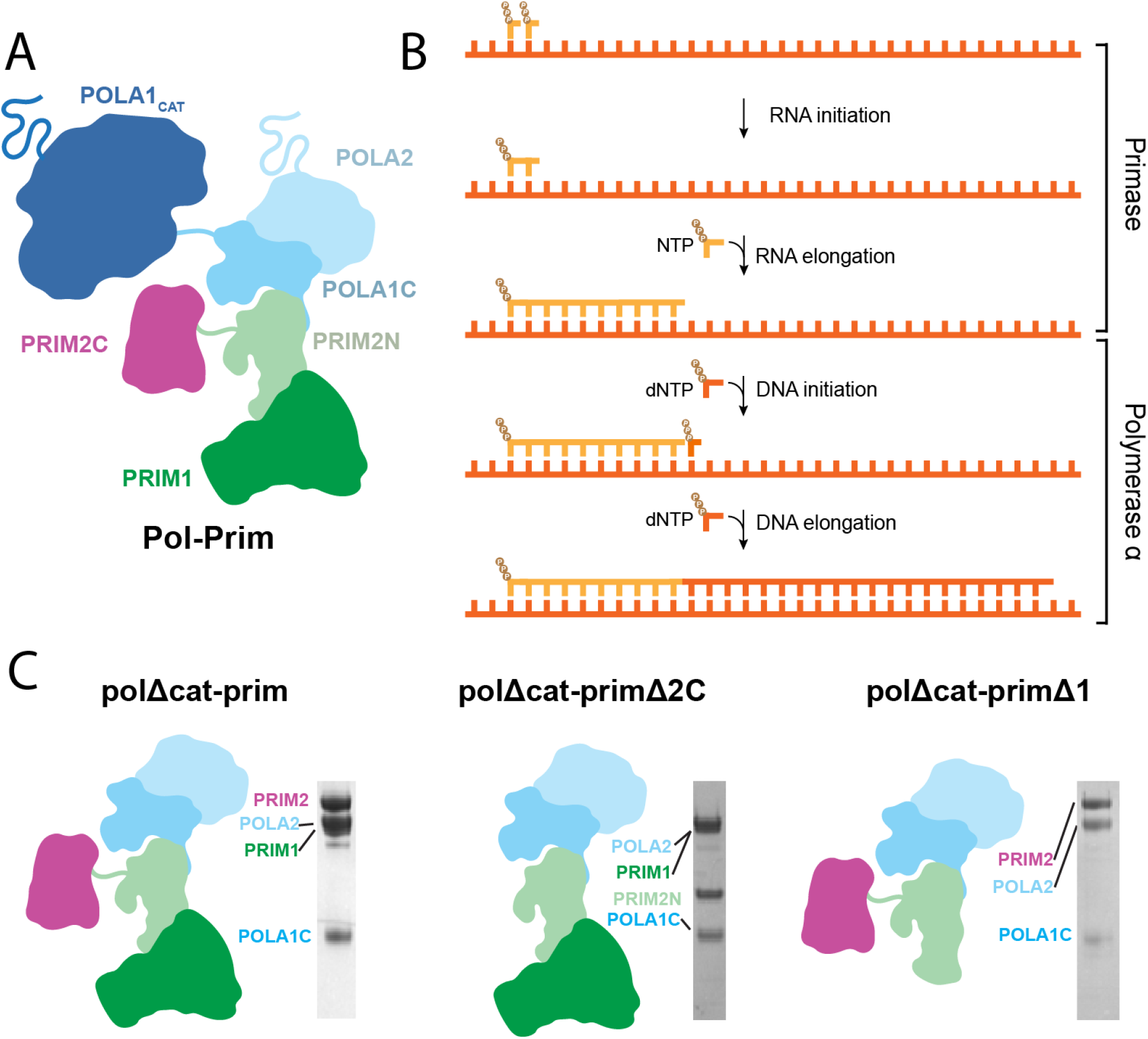
Schematic diagrams of primer synthesis and pol-prim structure. A) The pol-prim heterotetramer. B) Steps of primer synthesis. C Diagrams of pol-prim and subcomplexes used in this study alongside SDS-PAGE images of the purified proteins.

Primase consists of the PRIM1 (also known as p48, p49, PriS, and Pri1) catalytic subunit and the PRIM2 (a.k.a. p58, PriL, and Pri2) regulatory subunit.^12^ PRIM2 contains PRIM2N (residues 1-245), a domain that acts as a scaffold linking the primase subunits to the pol α subunits, and the flexibly attached PRIM2C domain (residues 265-455), which is involved in regulation of both RNA and DNA synthesis.^13–20^ Although the catalytic site for RNA synthesis is located within PRIM1, PRIM2C greatly stimulates catalytic activity and is required for both initiation of primer synthesis and regulation of the length of the primer.^21–23^ Polymerase α consists of the catalytic subunit POLA1 (a.k.a. p180) and the regulatory subunit POLA2 (a.k.a. p68, p70, and B subunit).^24,25^ POLA1 contains a catalytic domain (POLA1cat) and a C-terminal domain (POLA1C, residues 1265-1462) that acts as a scaffold between POLA2 and PRIM2N, while also connecting to POLA1cat through a flexible linker.^17,26,27^ Architecturally, pol-prim is organized into three modules: the tetramer core (POLA2/POLA1C/PRIM2N/PRIM1; a.k.a. pol-prim platform), POLA1cat, and PRIM2C.^28–30^ POLA1cat and PRIM2C are tethered to the tetramer core through ∼30 and ∼20 residue flexible linkers, respectively. The modularity of pol-prim allows for dynamic coordination of RNA and DNA primer catalysis as well as interfacing with other replication factors.^31–37^

RNA primer synthesis involves the coordinated action of PRIM1 and PRIM2C. PRIM1 alone possesses an ability to elongate pre-synthesized RNA primers in the absence of PRIM2C, but is otherwise incapable of initiating primers and has low affinity for oligonucleotide substrates on its own.^10,38,39^ PRIM2C has an inherently higher affinity for substrates, in particular for structures mimicking RNA primed templates that contain a 5’ triphosphate and 3’ template overhang.^28,38,39^ It is generally understood that initiation of primer synthesis requires a configuration in which PRIM1 and PRIM2C are aligned and together bind the ssDNA template, along with two incoming NTPs and catalytic metals before synthesis of the initial dinucleotide can occur.^20,40–43^ Formation of the first dinucleotide is known to be the rate-limiting step of primer synthesis.^10^

Once the dinucleotide is formed, the RNA primer is elongated by PRIM1 in a 5’-3’ direction to a length of 7-10 nucleotides. However, PRIM1 lacks the ability to regulate primer length without PRIM2C.^21,39^ Additionally, PRIM1 synthesis is distributive in nature, as synthesis of primers less than the unit length of 7 is common and is increased in the absence of PRIM2C.^10,39,40^ PRIM2C not only has higher affinity than PRIM1 for the substrate, but is also remains bound to the 5’ end of the primer, which plays an essential role in primer length regulation.^39,44,45^ From this it can be inferred that PRIM2C and PRIM1 become spatially separated as the primer length grows. Recent studies by our groups and others have also demonstrated that PRIM2C forms an interaction with POLA1cat during DNA initiation and likewise remains bound during DNA elongation, playing a critical role in handoff and DNA primer length regulation.^45–47^ In the prevailing model for RNA primer synthesis, a steric clash between PRIM2C and PRIM2N arises when the RNA primer reaches a length of 9 or 10 nucleotides, providing an upper limit for primer length.^20,28,30^ Our recent work has shown that steric clash with PRIM2C prevents POLA1cat from binding an RNA primer of <7 nucleotides, providing the lower bound of unit primer length as 7.^46^

Despite these advances in understanding that PRIM2C regulates primer synthesis, direct structural characterization of the priming complex during RNA catalysis is lacking. Here we report new data and present a mechanistic model for the initiation, elongation, and handoff of the RNA primer. By comparing binding affinities for RNA-primed template substrates of a variety of pol-prim constructs, we directly show that the majority of the substrate binding affinity of primase derives from the tight binding of PRIM2C. We also characterize structural transitions that occur as the RNA primer is initiated and elongated showing there is a mixture of states in all cases and that in the presence of substrate, the majority of molecules have PRIM2C but not PRIM1 bound. Our results reveal the molecular basis for the highly distributive nature of RNA primer synthesis and lead to a revised model for RNA primer synthesis that incorporates the high affinity of PRIM2C for substrate, the high-degree of flexibility of PRIM2C, and the preferential binding of RNA-primed templates by POLA1cat.

## Results

### Analysis of RNA priming by pol-prim requires specific construct design

The primase subunits of pol-prim (PRIM1, PRIM2) are responsible for the synthesis of the initial 7-10 nucleotide RNA primer. Moreover, the large POLA1cat domain, flexibly linked to the central tetramer core, is not directly involved in RNA primer synthesis. To focus our analysis of RNA priming we therefore utilized a truncated pol-prim subcomplex that lacks POLA1cat (PRIM1/PRIM2/POLA1C/POLA2, polΔcat-prim, Figure 1C), similar to that used for previous studies.^17,28,39,46^ The truncation of POLA1cat is advantageous for several reasons. Among these, the flexible nature and large size of POLA1cat would complicate measurements probing the conformational dynamics of PRIM1 and PRIM2C, which are the primary focus of this study. Deletion of POLA1cat is also valuable because it binds RNA-primed templates with higher affinity than PRIM1, so its absence ensures efficient generation of complexes relevant to RNA primer catalysis upon addition of relevant substrates.^38,39,46,48,49^ An additional important factor is that in the absence of nucleotide substrates, full length pol-prim exists primarily in a compact configuration (termed the Autoinhibitory conformation) in which POLA1cat forms physical interactions with and restricts the range of motion of both PRIM2C and the tetramer core.^28,45,46,50^ Pol-prim must drastically rearrange from this configuration before primer synthesis can occur as both the substrate binding site of PRIM2C and the DNA catalytic site of POLA1cat are occluded. Thus, removal of POLA1cat allows the analysis of the inherent conformational flexibility within the tetramer core as well as flexibility of PRIM2C relative to PRIM1.^27^

Recent structures of pol-prim in complex with the CST complex critical for telomere processing have established that CST acts as a scaffold for pol-prim that also restricts the flexibility and range of motion of the tetramer core and PRIM2C.^43,50,51^ In this study, we did not use either CST or other binding partners of pol-prim such as RPA that may influence the structure and dynamics of pol-prim because our goal was to establish the inherent degree of flexibility in polΔcat-prim. Our data also inform on the extent to which scaffolding may alter flexibility and how this may contribute to the mechanism of primer synthesis by pol-prim.

### The pol-prim tetramer core is conformationally heterogeneous

The flexible linkers connecting POLA1cat and PRIM2C to the tetramer core are critical to the structural changes in pol-prim that are required to initiate and synthesize the chimeric RNA-DNA primers.^28,29^ For RNA priming, the positioning of PRIM2C with respect to the tetramer core is key. The tetramer core has traditionally been viewed as a stable globular platform on which the primer is assembled.^28,30^ However, multiple structures of pol-prim that have emerged in the past few years reveal a significant degree of conformational heterogeneity within the core.^28,43,45–47,50,51^ Superposition of the tetramer core from these structures suggests there are hinges at the POLA1C/PRIM2N interface and within PRIM2N itself. Upon alignment to POLA1C and POLA2, which together form a rigid body,^24^ PRIM1 is seen to occupy a range of orientations that differ from each other by up to 35°, corresponding to a shift of ∼40 Å with respect to the center of PRIM1 (Fig 2A). Hence, in order to characterize the structural dynamics of PRIM2C relative to PRIM1, it is important to understand the conformational heterogeneity within the tetramer core.

**Figure 2.**
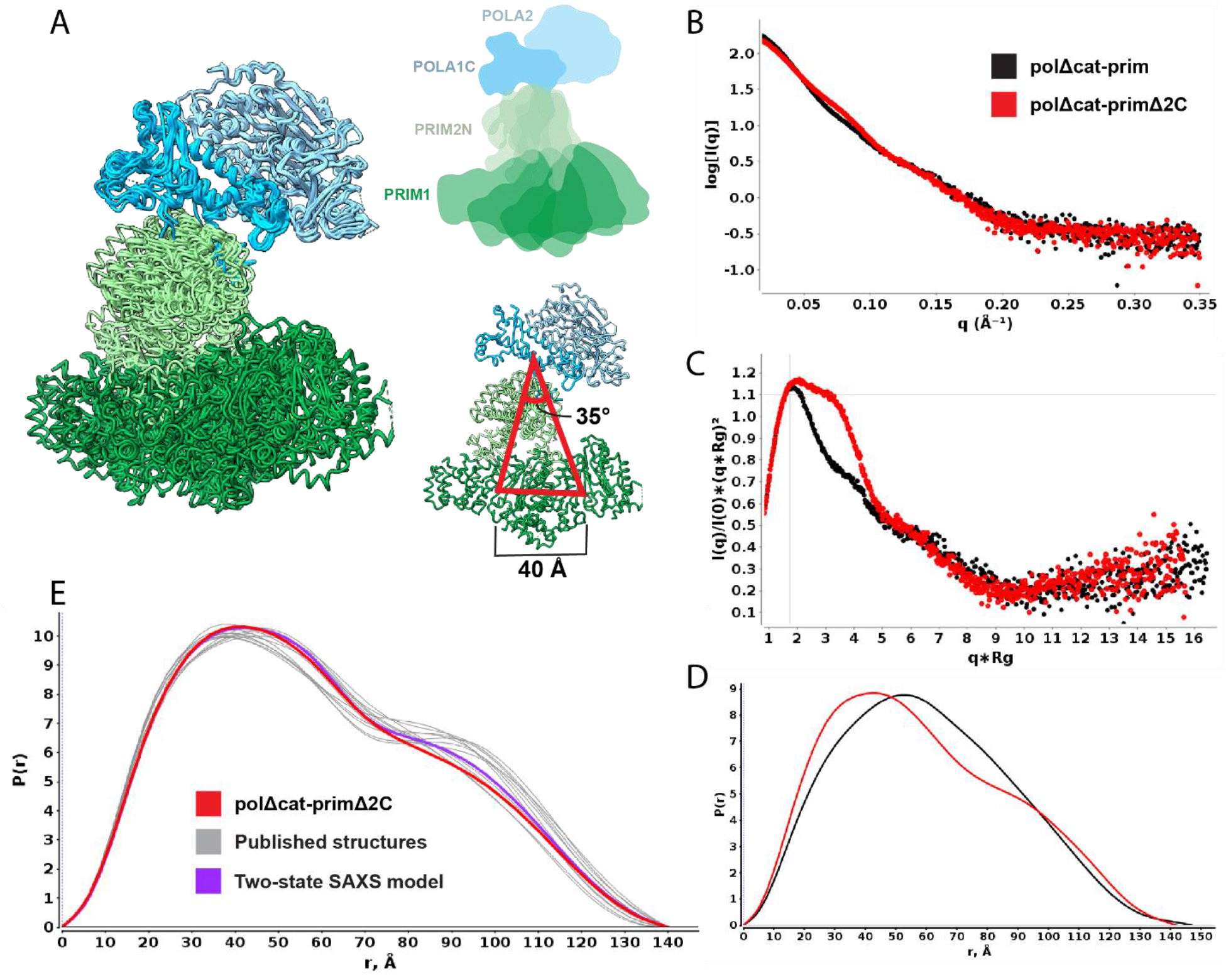
Flexibility of tetramer core in solution. A) Extracts from structures containing the tetramer core aligned to POLA1C/POLA2/PRIM2N (light blue), showing range of motion in PRIM2N/PRIM1 (green). Models are aligned to POLA1C/ POLA2. Comparison of SAXS data for free polΔcat-prim (black) and polΔcat-primΔ2C (red): B) log10 intensity plot; C) Normalized Kratky plot; D) Distance distribution. E) P(r) distribution of polΔcat-primΔ2C (red) overlaid with back-calculated distance distributions from coordinates extracted from available structures (gray) and two-state multi-FoXS model (purple).

To directly characterize the degree of flexibility within the tetramer core, we turned to small angle X-ray scattering (SAXS) analysis of a construct in which both POLA1cat and PRIM2C were removed so that just the tetramer core remains (polΔcat-primΔ2C, Figure 1C). This construct proved to be highly amenable to SAXS analysis and high quality, aggregation-free scattering data were acquired (Figure 2, Table 1). The Kratky plot of the SAXS data (Figure 2C) is characteristic of a multi-domain globular species and the shape of the corresponding P(r) distribution (Figure 2D) indicates the particle is dumbbell shaped, as expected based on the available pol-prim structures.

**Table 1.**
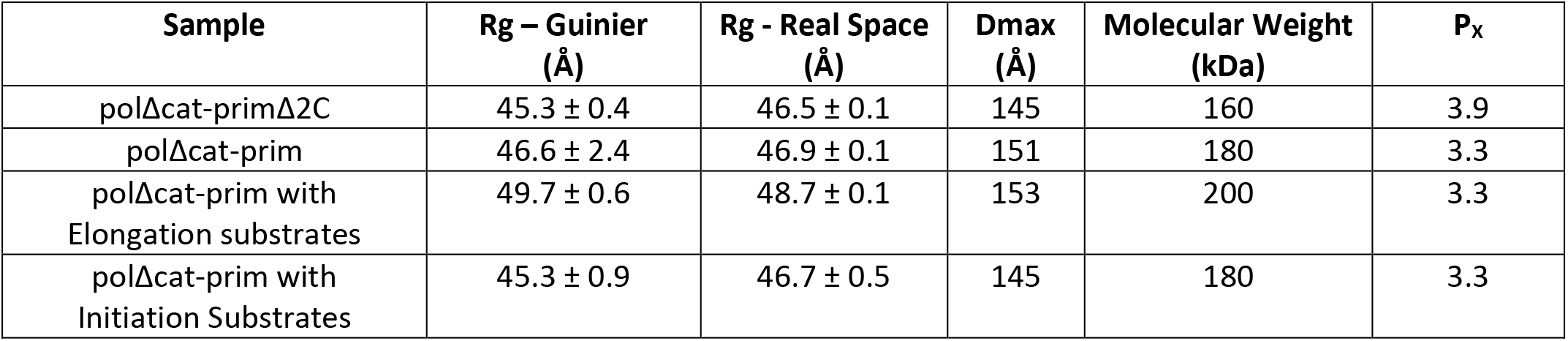
SAXS parameters.

To determine if there is a correspondence between the extent of heterogeneity in the tetramer core in published pol-prim structures and of polΔcat-primΔ2C in solution, we extracted the coordinates for the tetramer core from the ensemble of structures and back-calculated the corresponding SAXS curves (Figure 2A, E). The fits to the experimental data of single back-calculated scattering curves produced chi-squared values ranging from 4.9 to 11.4, indicating that none of the individual structures is a proper representation of the structure in solution. However, multi-state modeling of the SAXS data using Multi-FoXS with the ensemble of structures produced much better fits, and these could be futher improved by weighting the mixture of input structures.^52^ Interestingly, the best fit to the data using the simplest two state model was obtained for the two extremes of the ensemble of structures (chi-squared fit of 1.8) (Figure 2E). Overall, the conformational heterogeneity observed in the available pol-prim structures and the SAXS data are in good agreement that the tetramer core is not a fixed structural platform but rather samples a continuous mixture of conformations in solution, consistent with our cryoEM analysis of full-length pol-prim.^46^

### PRIM2C is a primary contributor to pol-prim configurational flexibility

Variation in the orientation of PRIM2C with respect to the tetramer core has been inferred to play a critical role in facilitating primer synthesis and regulating the length of both the RNA and DNA portions of the primer.^21,43,45–47,54,55^ It has long been established that PRIM2C is attached to the tetramer core of pol-prim by a flexible linker.^17,20,30^ Among the eleven available pol-prim structures in which PRIM2C is visible, eight occupy a configuration roughly similar to the autoinhibited state, in which PRIM2C is located near the PRIM2N and POLA1C interface and constrained by direct contacts with POLA1cat.^28,46,47,50^ Three additional orientations are observed in recent structures of pol-prim in the presence of single-stranded DNA or an RNA-primed template.^43,45^ Although there is some variation observed in the available structures, knowledge of the nature and degree of the PRIM2C flexibility remains incomplete.

To obtain direct insight into the extent of configurational variability of PRIM2C, we collected SAXS data for the tetramer core with PRIM2C attached (polΔcat-prim). These data, when compared to the data described above for the tetramer core construct lacking PRIM2C (polΔcat-primΔ2C), provide critical insights into the flexibility of PRIM2C (Figure 2B-D, Table 1). As was the case for for polΔcat-primΔ2C, polΔcat-prim proved to be highly amenable to SAXS analysis and high quality, aggregation-free scattering data were acquired. The Kratky plot of the SAXS data indicates that polΔcat-prim is a mostly globular particle but with significant interdomain flexibility (Figure 2C), showing directly what was previously inferred, i.e. that the attachment through a ∼20-residue flexible linker results in PRIM2C occupying a range of relative orientations with respect to the tetramer core. Significant inter-domain flexibility is also reflected in the substantially lower value of the Porod Exponent (P_X_) for polΔcat-prim (∼3.3) relative to the tetramer core alone (polΔcat-primΔ2C, 3.9). [Unlike the conformational heterogeneity *within* the tetramer core, the substantial changes in the orientation of PRIM2C relative to the tetramer core have a substantial impact on the surface-area to volume ratio and correspondingly result in the lower value of P_x_.] In addition to these parameters, the Mid-Q regions of the intensity and Kratky plots of the two constructs are clearly different (Fig 2B, C), as are the corresponding P(r) distributions (Figure 2D). These parameters reflect a cylindrical shape for polΔcat-primΔ2C, whereas the attachment of PRIM2C in polΔcat-prim results in a more dumbbell-like shape. Although it samples a variety of orientations, the SAXS data suggest that on average PRIM2C is located somewhat centrally between the POLA1C/POLA2 lobe and the PRIM1/PRIM2N lobe of the tetramer core.

In order to better understand the distribution of configurations of PRIM2C with respect to the tetramer core, we analyzed the structure of polΔcat-prim by electron microscopy (Figure S2A). Negative stain EM images showed clear particles with the expected dumbbell-like shape of polΔcat-prim. Excellent density was observed in 2D class averages for the bulk of the tetramer core, albeit with somewhat blurred density for PRIM1. The observation of blurred density for PRIM1 corresponds well to the flexibility of PRIM1 with respect to the rest of the tetramer core arising from the hinge within PRIM2N. In contrast, the density for PRIM2C was highly blurred and 3D reconstructions lacked consistent density for PRIM2C. A small, disconnected segment of density was visible in some reconstructions, but these could not be assigned to PRIM2C with confidence. The lack of density for PRIM2C indicates that it occupies a range of configurations in solution rather than a limited number of discrete states, consistent with our SAXS analysis.

To improve the resolution of our negative stain EM images, we turned to crosslinking polΔcat-prim with BS3. In 2D class averages strong density was observed for the tetramer core (including PRIM1) from a variety of viewing angles (Figure 4A, Figure S3). In both 2D class averages and 3D reconstructions, the conformation of the tetramer core was noticeably more homogenous than in the non-crosslinked sample and the characteristic S-shaped bilobal architecture was clearly evident. Moreover, in 2D class averages PRIM2C was readily visible and occupied a continuous range of configurations spanning from near PRIM1 to the opposing orientation near the interface of PRIM2N and POLA1C as observed in the structures of the Autoinhibitory state.^28,45–47,50^ 3D reconstruction and classification yielded two primary classes, one in which PRIM2C was centrally located as in the SAXS based models, and the other in which PRIM2C was located near POLA1C as in structures of the Autoinhibitory state (Figure 4, S4).^28,45,46,50^ These are presumably two well-populated orientations from within the ensemble of configurations accessible to PRIM2C that have been selected for by crosslinking.

**Figure 3.**
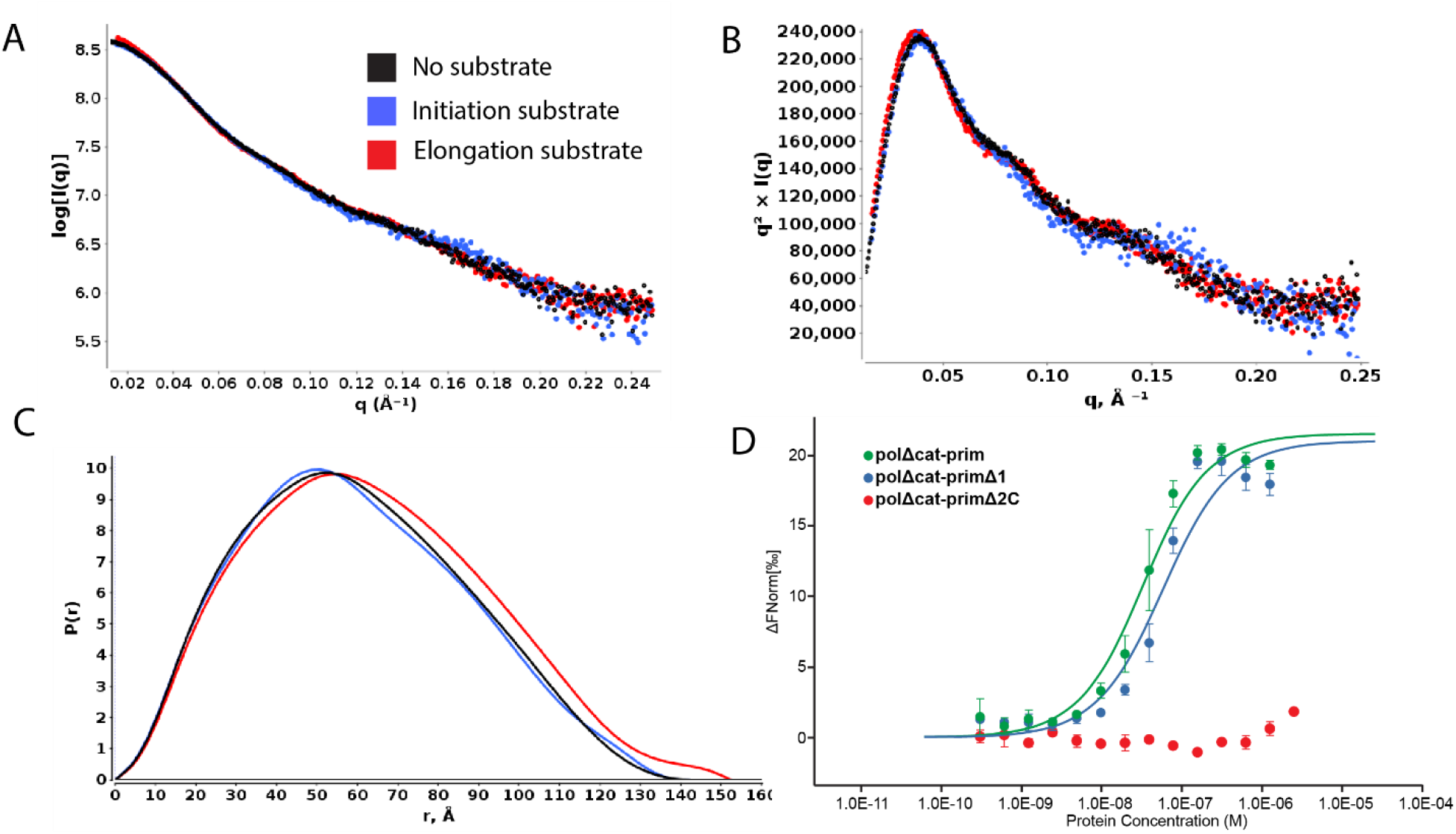
RNA elongation substrate engagement in solution. Comparison of SAXS data for free polΔcat-prim (black) and polΔcat-prim in the presence of an RNA elongation substrate (red), and substrate and co-factors required for initiation (blue): A) log10 intensity plot; B) normalized Kratky plot; C) distance distribution. D) MST dose-response curve measuring RNA elongation substrate binding of polΔcat-prim (green), polΔcat-primΔ1 (blue), and polΔcat-primΔ2C (red).

**Figure 4.**
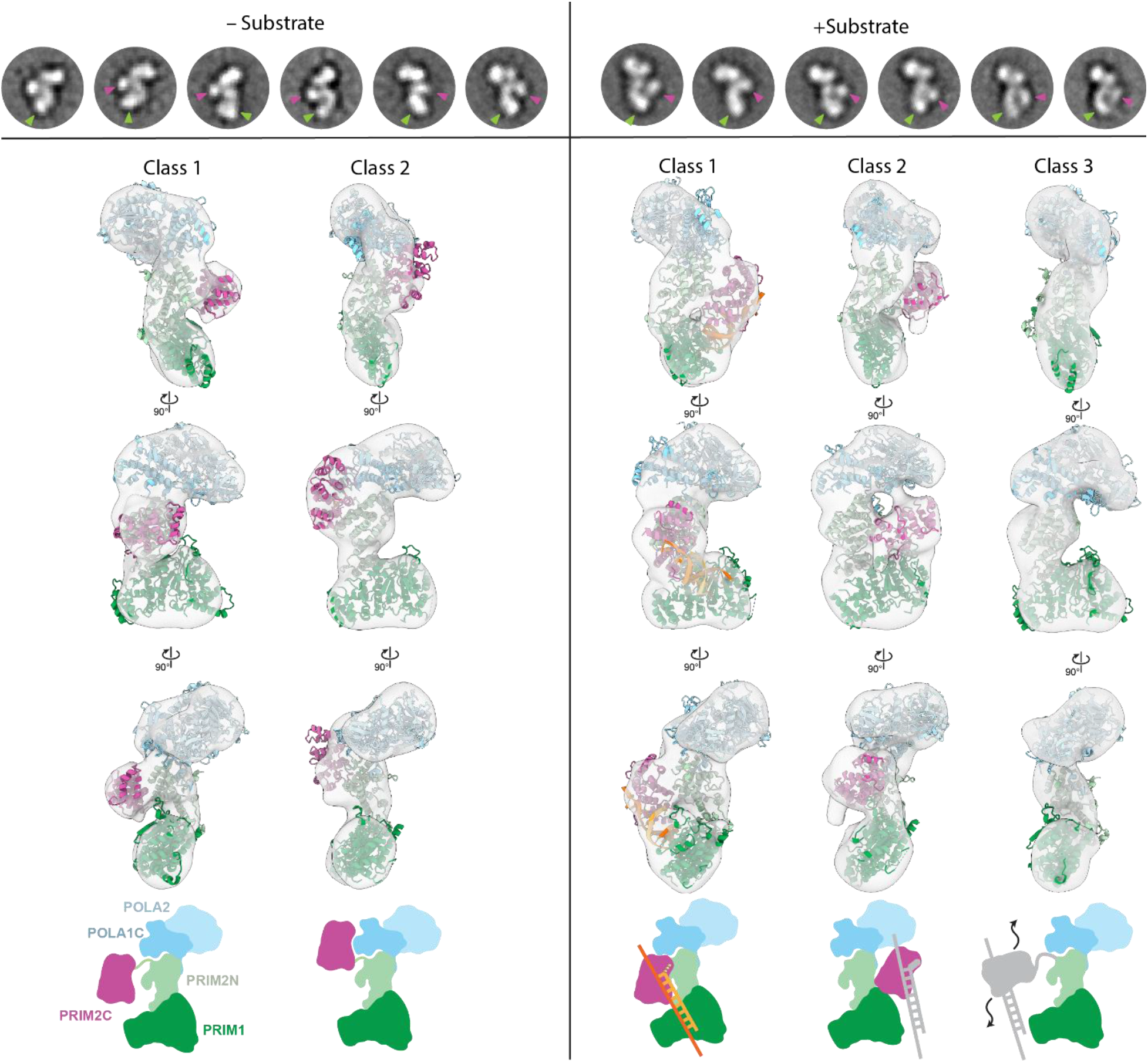
PRIM2C orientation and heterogeneity visualized by Negative Stain EM. Top row-selected 2D class averages of prim +/- elongation substrate with PRIM2C (purple) and PRIM1 (green) density highlighted. Comparison of 3D reconstructions without (left) and with (right) elongation substrate. Atomic models rigid-body docked into the density is shown for polΔcat-prim. The +substrate class 1 model corresponds to the elongation model shown in Figure S5. Bottom row-schematic diagrams drawn to correspond to atomic models to which they align.

### PRIM2C does not frequently sample the catalytically active RNA initiation configuration

The changing orientation of PRIM2C is key to the progression through the multiple stages of primer synthesis. To better understand these changes, we set out to characterize the structural flexibility of polΔcat-prim under conditions required for RNA primer initiation and when bound to an RNA-primed template. We first sought to determine if a SAXS analysis would be sufficiently sensitive to detect the anticipated differences in the structure by carrying out a series of simulations. We generated two structural models of polΔcat-prim: one in the configuration required for initiating RNA primer synthesis and the other in the configuration when bound to an 8mer RNA primed template elongation substrate (Figure S5A,E). The model for initiation requires that PRIM2C is located over the active site of PRIM1 and engaged in binding nucleotide. In generating our model for the complex with the elongation substrate, PRIM2C had to be moved away from PRIM1 as PRIM2C remains bound to the 5’ triphosphate moiety of the nascent RNA primer as PRIM1 extends the 3’ end.^28,39^ Our model of polΔcat-prim in an actively elongating configuration is notably different from a recent structure of yeast pol-prim in the presence of an RNA-primed template in which PRIM2C is associated with the primer but the 3’ end of the growing primer is ∼27 Å away from the PRIM1 active site in a catalytically inactive configuration (Figure S5G).^45^

The polΔcat-prim models for initiation and elongation of RNA synthesis were used to back-calculate SAXS data to determine how much they differed. To have a point of reference, these were overlayed on the experimental scattering data of polΔcat-prim alone (Figure S5C,F). The back-calculated scattering profiles and distance distributions for the models confirmed that the changing orientation of PRIM2C with respect to the tetramer core during primer initiation and elongation has a significant impact on the corresponding SAXS parameters. Based on the prediction that the magnitude of shape change in the presence of substrate would be detectable by SAXS, we designed two experiments to acquire SAXS data corresponding to these two key points in the trajectory of RNA primer synthesis.

The first experiment involved acquiring SAXS data for polΔcat-prim in the presence of a large excess of all components required for initiation: template single-stranded DNA, NTPs (in the form of the non-hydrolyzable nucleotide analogue GMPCPP), catalytic metal Mn^2+^ (Table 1). Large excesses of substrate and the co-factors were included because all of these are bound with only modest affinities in the low micromolar to high nanomolar range. Comparing the SAXS results to those for polΔcat-prim alone shows only a modest shift in the Mid-Q region of the log10 intensity and the Kratky plots (Figure 3A, B). Moreover, the P(r) distribution was mostly unchanged relative to the free protein (Figure 3C). These observations suggest that the presence of the substrate and co-factors needed for initiating RNA priming does not drive pol-prim to highly populate the configuration required for catalysis. Based on a series of simulations to determine the precision of differences between two experimental SAXS curves, we were able to estimate that only a small proportion of particles (∼10%) adopt the configuration required for initiating RNA synthesis even in the presence of excess template and co-factors. Initiation of RNA synthesis is understood to be the rate limiting step of primer synthesis,^10^ and from the ensemble of evidence now available we can propose a molecular mechanism to explain why: for RNA priming to initiate, pol-prim must not only sample the rare configuration in which both PRIM2C and PRIM1 are aligned but both must also have the initiation substrates and their requisite co-factors bound.

### PRIM2C flexibility is retained during RNA elongation

The second set of experiments on polΔcat-prim were performed with an 8mer RNA elongation substrate (5’ tri-phosphorylated 8mer RNA annealed to single-stranded DNA), designed to characterize PRIM2C configurational flexibility during RNA elongation of the primer strand. Surprisingly, examination of the SAXS intensity and Kratky plots revealed they were quite similar to those observed for the free protein (Figure 3A,B) even though there was a distinct change in size. Assurance that the complex was formed was provided by the molecular weight estimate, Rg, and Dmax, which all indicated that the ∼14 kDa RNA/DNA duplex substrate was bound (Table 1). The shift to the right in the P(r) distribution was also consistent with an increase in the overall size of the complex (Figure 3C).

To directly verify that the substrate was indeed bound and that both PRIM1 and PRIM2C were engaged, we carried out a series of substrate binding experiments using microscale thermophoresis (MST). To further dissect the relative contributions of PRIM1 and PRIM2C to substrate binding, affinities were measured for polΔcat-prim, the tetramer core construct (polΔcat-primΔ2C), and a third construct that deletes PRIM1 but retains PRIM2C (polΔcat-primΔ1) (Figure 1C). The substrate for these experiments was a 9mer RNA-primed template, designed specifically to maximize the stability of the duplex (Figure 3D, Table 2). As anticipated based on previous studies,^39^ we observed very tight binding of this substrate by polΔcat-prim (K_d_ = 28 ± 7 nM). The deletion of PRIM1 resulted in a two-fold decrease in binding affinity (K_d_ = 53 ± 18 nM), consistent with previously reported affinities for PRIM2C alone.^39^ In contrast, truncation of PRIM2C resulted in a significant decrease in affinity, below the limit of detection of the assay. Our data for polΔcat-prim show that PRIM2C is the primary contributor to substrate binding during RNA priming as suggest by previous studies of isolated PRIM2C and isolated primase (PRIM1/PRIM2).^20,38,39^

**Table 2.**
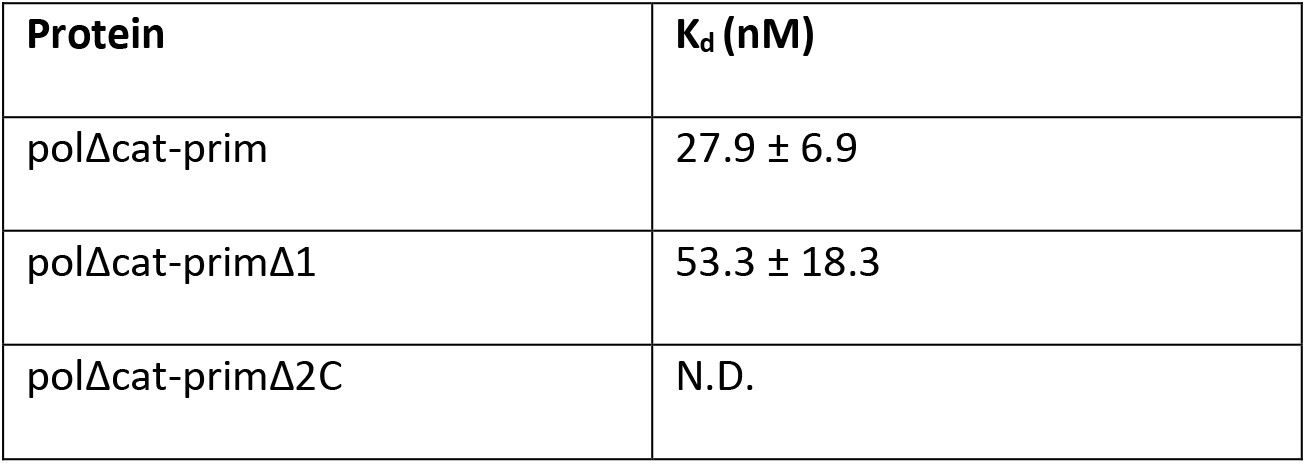
Estimated Dissociation Constants.

To better understand the distribution of configurations of polΔcat-prim with the elongation substrate bound, we again turned to EM imaging and compared the data to that of free protein. As expected, there was no difference in the extent of conformational variability within the tetramer core (Figure S2B). There was however somewhat better density for both PRIM1 and PRIM2C in both 2D class averages and a 3D reconstruction, suggesting reduced configurational variability, as might be anticipated when a ligand is bound. Nevertheless, in most 2D class averages, the density for PRIM2C was blurred and in a similar position to that seen in the absence of substrate. However, in a small subset of classes PRIM2C appeared to be oriented near to PRIM1, an orientation that was not obviously apparent in the absence of substrate. Overall, although there was improved density in the class averages and 3D reconstruction, a high degree of variability in the positioning of both PRIM2C and PRIM1 was still evident even when the elongation substrate is bound.

How does one rationalize that the differences between the free protein and elongation complex are so small? The key factor is the low affinity of PRIM1 for substrates relative to the very high affinity of PRM2C, which indicates that for the majority of the polΔcat-prim molecules PRIM2C will be engaged but PRIM1 will not. As a result, PRIM2C can move away from its alignment over PRIM1, maintaining a similar level of configurational freedom to what is observed for free polΔcat-prim.

### Configurational freedom of PRIM2C and substrate dissociation from PRIM1 are key features during RNA priming

RNA elongation requires population of a pol-prim configuration in which the 3’ end of the growing primer is engaged in the PRIM1 active site, while PRIM2C remains tightly bound to the 5’ end of the primer. Although the EM data acquired without cross-linking provided some evidence of the orientation of PRIM2C in an active configuration, sorting of particles into discrete classes was hampered by the high degree of ambiguity due to PRIM1 and PRIM2C flexibility. To characterize the configurational flexibility of PRIM2C during RNA elongation in more detail, we crosslinked the complex of polΔcat-prim with the RNA elongation substrate using BS3. The conditions for crosslinking were optimized by following the production of crosslinked products by SDS-PAGE and by comparing the results with and without substrate (Figure S6). A series of crosslinked products is observed on the gel in the absence of substrate corresponding to various combinations of the subunits, but with the four-subunit product being a predominant species. With the addition of substrate, these bands shifted to slightly higher molecular weight, consistent with successful crosslinking of the substrate to polΔcat-prim.

Analysis of the negative stain EM data (Figure S4B, S7) revealed the overall architecture of polΔcat-prim was retained when bound to the elongation substrate with the tetramer core adopting the conformation observed previously (Figure 4). In 2D class averages, a shift in the proportion of classes was observed in which PRIM2C and PRIM1 were aligned and formed continuous density, indicating enrichment of particles with a configuration in which both PRIM1 and PRIM2C are bound to the elongation substrate (Figures 4, S7). However, a subset of 2D classes were present in which PRIM2C was not aligned over PRIM1 and instead occupied a central location, as expected if PRIM1 was not bound to the substrate. 3D reconstruction yielded three primary classes (Figures 4, S4B). In class 1 (36%), clear density was visible for PRIM2C and formed continuous density with PRIM1, consistent with both PRIM1 and PRIM2C being bound to substrate. Interestingly, insertion of our model of polΔcat-prim bound to an RNA elongation substrate into the density revealed an excellent overlap in the position of PRIM2C. In class 2 (34%), density for PRIM2C was observed, but was shifted away from the tetramer core while still exhibiting some continuous density with PRIM1, though to a lesser extent than in class 1. This class has additional density protruding from PRIM2C, which we attribute to the substrate being bound to PRIM2C but not PRIM1. A recently published structure of pol-prim in complex with an RNA-primed template reveals PRIM2C bound to the substrate but positioned away from the active site of PRIM1 (Figure S5G).^45^ While there is considerable uncertainty about the author’s assignment of this structure as an RNA synthesis state because the primer lacks the critical 5’ tri-phosphate that is key to PRIM2C interaction and required for RNA priming, we note a general similarity to class 2 of our 3D reconstructions. In class 3 (30%), density for the tetramer core was very similar to the density observed in the absence of substrate, but density for PRIM2C was not visible. We believe that like class 2, the substrate is not bound to PRIM1, but the density of PRIM2C is absent because there is greater degree of PRIM2C configurational heterogeneity of the particles contributing to this class.

Interestingly, the location of PRIM2C in both class 1 with substrate and class 1 without substrate is very similar. However, comparison of the density connecting PRIM2C and PRIM1 reveals a substantial difference, especially at lower contour levels (Figure 5, S8). In the absence of substrate, a clear separation between PRIM2C and PRIM1 is visible corresponding to a species in which PRIM2C is disengaged from PRIM1. In the presence of the elongation substrate, clear tubular density is visible connecting PRIM2C and PRIM1, nicely corresponding to the location of the RNA/DNA duplex in the atomic model of the elongation complex.

**Figure 5.**
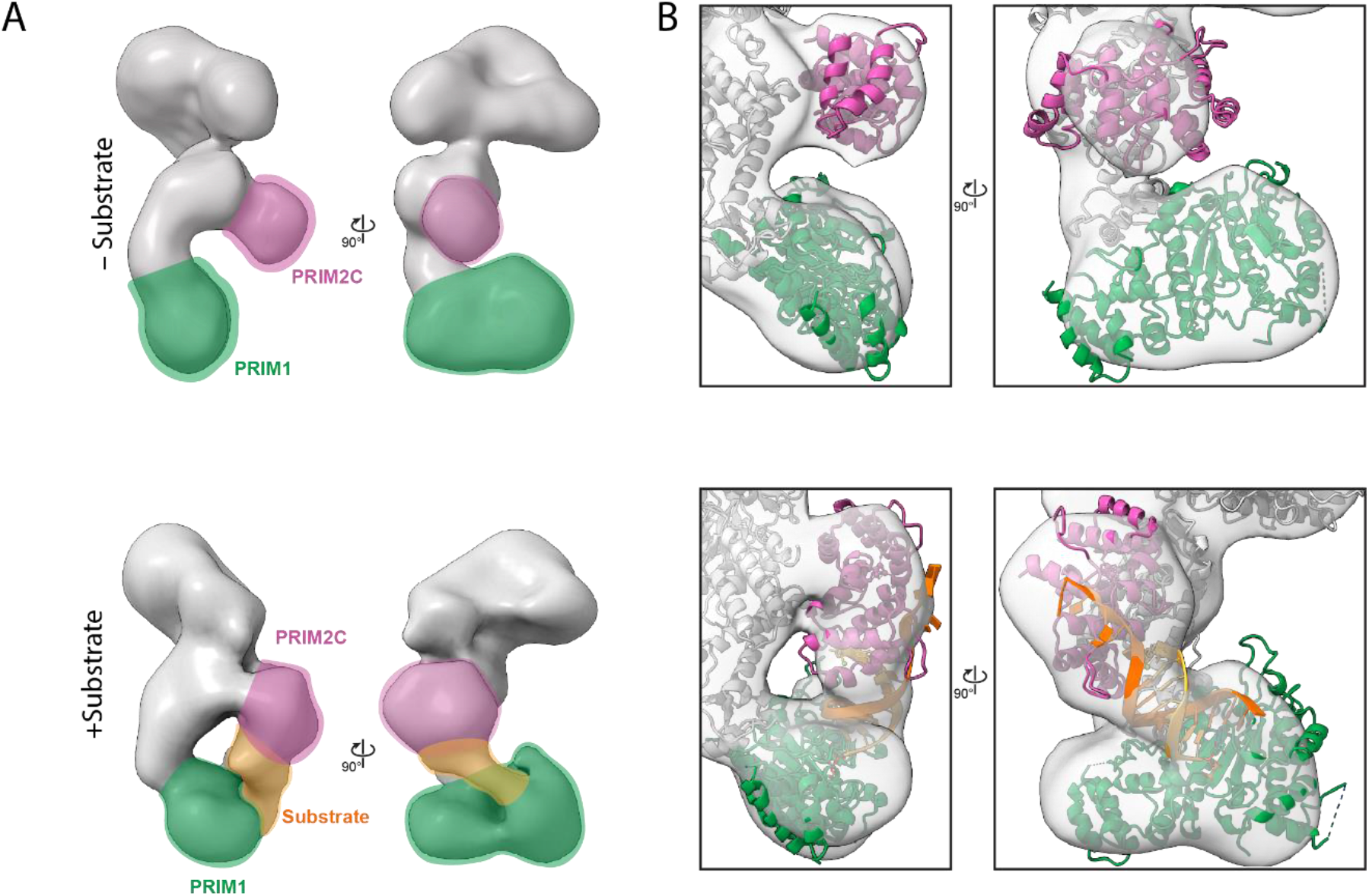
3D map of polΔcat-prim and the complex with the RNA elongation substrate. A) 3D reconstruction of class 1 without (top) and with substrate (bottom). The density for PRIM2C (purple) PRIM1 (green), and substrate (orange) are highlighted. B) Position of PRIM2C and PRIM1 from atomic models fit into the maps obtained without and with substrate.

Overall, a significant degree of configurational variability in PRIM2C is evident even after crosslinking. In the absence of substrate, PRIM2C appears to occupy a continuous range of configurations, sampling many possible orientations. In the presence of substrate, there remains a significant range of configurations of PRIM2C, but the origin is different: it is the dissociation of the substrate from PRIM1 that allows a higher than anticipated degree of configurational freedom for PRIM2C.

## Discussion

The basic steps of primer synthesis by pol-prim involves formation of the initial dinucleotide by the concerted action of PRIM1 and PRIM2C, then elongation to 7-10 nucleotides by PRIM1 before the RNA primer is handed off (transferred intramolecularly) to POLA1cat.^9–11,56^ PRIM2C has been shown to play roles in RNA initiation, elongation, length regulation, and handoff.^21,38^ However, the structural basis for how PRIM2C functions is incomplete, having relied heavily on inference from biochemical experiments and structures of PRIM2C alone or of larger pol-prim constructs in the absence of substrates.^20,28–30^

Our investigation of polΔcat-prim provides key structural context for assessing proposed biochemical models of RNA primer synthesis. In particular, we directly show that PRIM2C has inherent configurational flexibility, visualizing a wide range of orientations relative to the tetramer core. In addition to PRIM2C flexibility, we expand upon recent studies to directly characterize the conformational heterogeneity of PRIM1 in the tetramer core.^20,28,43,45,46^ These high degrees of flexibility within the primase subunits provide the adaptive architecture needed to accommodate the multivalent and semi-independent substrate binding by PRIM2C and PRIM1 that is characteristic of RNA primer initiation and elongation.

Our binding studies systematically tested the proposal that PRIM2C is the primary contributor to substrate binding affinity during RNA priming.^38,39^ Our structural studies revealed the very low population of pol-prim configurations in which PRIM1 and PRIM2C are aligned for synthesis of the initial dinucleotide even when the template and co-factors are present in large excess. For the complex with an RNA elongation substrate, crosslinking enabled visualization of three primary states: one in which PRIM2C and PRIM1 are both bound to the primer in a configuration competent for RNA elongation, the second in which PRIM2C is visible and bound to substrate but has moved away from PRIM1, and the third in which PRIM2C is not visible and is structurally independent from PRIM1 and the rest of the tetramer core. This degree of configurational variability of PRIM2C even in the presence of substrate: (1) provides evidence in support of the proposal that PRIM1 dissociates from the primer during synthesis, (2) directly correlates with the distributive nature of RNA elongation by primase, and (3) further contextualizes structures of pol-prim in the presence of an RNA-primed template.^20,45^

The large difference in the affinity of PRIM1 and PRIM2C for substrates is consistent with the apparent dominance of PRIM2C-bound/PRIM1-unbound particles and has important implications for the mechanism of primer synthesis. First, the asymmetry in the interaction with the substate provides a structural basis for the distributive (rather than processive) nature of primer synthesis by PRIM1 that has been established by previous biochemical analyses.^9,10,21^ Those studies showed that PRIM1 synthesizes abortive primer products (di- or trinucleotides) in abundance in both the presence and absence of PRIM2C, while the presence of PRIM2C results in an increase in unit length 7-10-nucleotide primers. Thus, PRIM2C acts as a processivity factor, decreasing the dissociation of short products before synthesis of the RNA primer can be completed.^20,28^ PRIM1 also elongates pre-made primers to lengths of up to 40 nucleotides in an unregulated fashion in the absence of PRIM2C, while PRIM1/PRIM2 synthesizes primers either of defined length (7-10 nucleotides) or as multiples of 7-10 nucleotides.^21,39^ Hence, PRIM2C also acts as a length regulator, influencing both the maximum and minimum primer length.

Our EM reconstruction in which PRIM2C and PRIM1 both engage an RNA-primed template substrate is the first structure of pol-prim bound to substrate in a configuration competent for RNA synthesis. It also supports models of primer synthesis in which PRIM2C remains engaged throughout. In one such model, the upper bound for primer length is attributed to a steric clash between PRIM2N and PRIM2C at a primer length of ∼10 nucleotides.^28^ This model is consistent with the close proximity of the PRIM2C lobe and the tetramer core in our reconstruction of the complex with the RNA elongation substrate. Our recent studies of Xenopus pol-prim provide further insight, showing that the minimal length of 7 nucleotides of RNA is due to the inability of POLA1cat to bind primers shorter than 7 nucleotides as a result of steric occlusion by PRIM2C.^46^

Our results provide the basis for a unified model of RNA priming by pol-prim, which is consistent with all available biochemical and structural data and is outlined in schematic form in Figure 6. The initiation of primer synthesis requires alignment of PRIM2C and PRIM1 in a catalytically competent configuration as well as binding of the two nucleotides, the catalytic metals and the ssDNA template. The sparse population of pol-prim configurations with PRIM1 and PRIM2C aligned combined with the low binding affinities for the template, nucleotides and metals makes initiation the rate-limiting step of primer synthesis. After formation of the initial dinucleotide, PRIM1 extends the RNA primer in a distributive manner due to dissociation of the nascent primer from the PRIM1 active site, the result of its low affinity for nucleotide substrates. PRIM2C functions as a processivity factor, remaining bound to the 5’ end of the primer and thereby facilitating PRIM1 re-association with the 3’ end of the primer. The lower bound for PRIM1 extension is 7 nucleotides because, as we have shown elsewhere, this is the minimum primer length necessary binding by POLA1cat.^46^ Importantly, POLA1cat has also been demonstrated to exhibit high affinity for substrate in the nanomolar range, several orders of magnitude higher than the affinity of PRIM1 for RNA-primed template we observed.^25,46,48,57,58^ Thus, after the primer has reached 7 nucleotides, competition for the primer between PRIM1 and POLA1cat occurs in which POLA1cat is highly favored. In this model, the low affinity of PRIM1 for substrate means that the primed template is inherently available for transfer to POLA1cat. An additional contribution may come from an inherent orientational preference of PolA1cat that strategically positions it to quickly bind the 3’ end of the RNA primer after PRIM1 dissociation.^45^ Extension of the nascent primer by PRIM1 can occur up to 10 nucleotides if PRIM1 dissociation does not occur, after which steric clash between PRIM2C and the tetramer core greatly suppresses further elongation.

**Figure 6.**
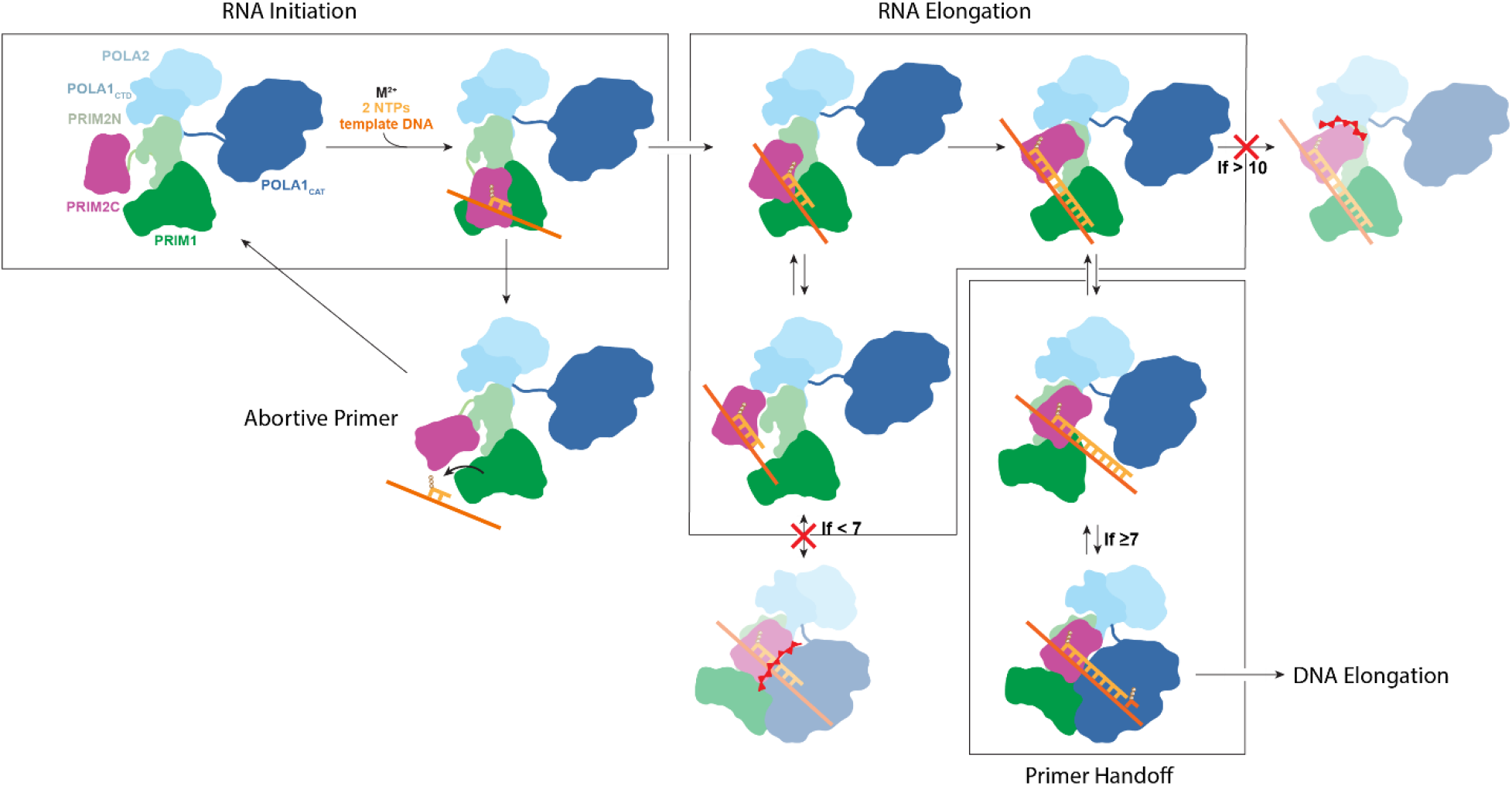
Model of RNA primer synthesis. Initiation of RNA synthesis requires PRIM2C and PRIM1 to be aligned with template, catalytic metals and nucleotides bound to enable synthesis of the initial dinucleotide. The dinucleotide is elongated by PRIM1 while PRIM2C remains bound to the 5’ triphosphate and progressively moves away from the active site of PRIM1 as the primer grows. The primed template dissociates from PRIM1 during primer synthesis due to its low affinity for the substrate. PRIM2C acts as a processivity factor by remaining bound to the primed template, which facilitates re-association with PRIM1. The primer can continue to lengthen until a steric clash between PRIM2C and PRIM2N prevents further lengthening when the primer reaches 10 nucleotides. The ready dissociation of PRIM1 facilitates handoff of the primer to POLA1cat for synthesis of the DNA portion of the primer, although handoff cannot occur until the primer reaches the minimum of 7 nucleotides required for POL1cat binding primed template with high affinity.

Although pol-prim is capable of performing all steps of primer synthesis in isolation, it is important to take into consideration the many interaction partners that recruit and modulate pol-prim activity at the replication fork.^33,37,43,50,51,59–61^ Recent structures of pol-prim bound to CST suggest that: (1) CST acts as a pol-prim recruitment factor to telomeres; (2) CST can promote a pol-prim configuration in which the PRIM1 and PRIM2C are aligned for primer initiation while allowing for PRIM2C flexibility such that the PRIM1 and PRIM2C could spatially separate as the primer elongates.^43,50^ At replication forks, RPA may fulfill a similar function and facilitate positioning of PRIM2C and PRIM1 for primer initiation while strategically orienting POLA1cat for primer handoff. How such interactions are maintained or affected by additional components of the replisome remains poorly understood. Future studies to investigate the influences of these factors on the structural organization and flexibility of pol-prim throughout the various steps of chimeric primer synthesis will provide deeper insight into how this dynamic molecular machine functions in its various cellular contexts.

## Methods

### Cloning and Expression

The gene encoding FL-PRIM1 was cloned into a pBG100 expression vector while the gene encoding FL-PRIM2 was cloned into a pETduet co-expression vector. POLA2 (residues 155-598) and POLA1C (residues 1265-1462) were cloned into a pET-duet expression vector encoding a 6xHis TEV cleavable tag. The disordered N-terminal region (residues 1-155) was truncated to facilitate expression, purification, and biophysical analysis. To generate a PRIM2ΔC construct, the N-terminal domain PRIM2N (residues 1-265) was subcloned into a modified pETduet expression vector. To generate a polΔcat-primΔ1 construct, a separate tri-gene construct was generated which encoded POLA2 (residues 155-598), POLA1C (residues 1265-1462), and FL-PRIM2, which was then cloned into a pRSF-DUET vector.

PRIM1/PRIM2 and POLA1C/POLA2 were expressed separately in Escherichia coli BL21 RIL cells. PRIM1/PRIM2 cultures were grown at 37°C in LB medium supplemented with 25 mg/L chloramphenicol, 100 mg/L ampilicin, and 30mg/L kanamycin while POLA1C/POLA2 cultures were supplemented with chloramphenicol and ampicillin only. Upon reaching mid-log phase, cultures were cooled to 16 °C and PRIM1/PRIM2 cultures were supplemented with 100 mg/L ammonium iron (III) citrate and 100 mg/L iron (II) sulfate. Overexpression was then induced by addition of IPTG to a final concentration of 0.5 mM. After overnight incubation, cells were harvested by centrifugation and pellets stored at -80 °C.

### Protein Purification

PRIM1/PRIM2 and POLA1C/POLA2 cell pellets were thawed in room temperature water and resuspended in Lysis Buffer (50 mM sodium phosphate pH 7.5, 500 mM NaCl, 10% [v/v] glycerol, 1 mM TCEP, 0.1 g/50mL DNAse, 1 g/50mL lysoszyme, and 1 complete EDTA-free protease inhibitor cocktail tablet [Roche] per 50 mL). The pellets were co-lysed using a Dounce homegenizer followed by gentle sonication. The lysate was then centrifuged at 50,000 g for 1 hr to pellet cell debris. The supernatant was then filtered using a 0.45 micron filter before application to a Ni-NTA column on a FPLC (Biorad). The column was washed with 10 column volumes (CV) of 10 % Nickel Buffer A (Lysis buffer without DNAse, lysozyme, and protease inhibitor) 90% Nickel Buffer B (Nickel Buffer A + 1M imidazole), followed by protein elution at 60% Nickel Buffer B. Fractions containing the protein of interest were pooled, H3C and TEV protease were added for His-tag cleavage, and dialyzed overnight at 4 °C in dialysis buffer (20 mM HEPES pH 7.5, 200 mM NaCl, 5% glycerol, 1 mM TCEP) using 10 kDA MWCO dialysis tubing. The following morning, the dialyzed solution was diluted 1:1 using Dilution buffer (20 mM HEPES pH 7.5, 5% glycerol, 1 mM TCEP) and filtered using a 0.45 micron filter before application to a Heparin column pre-equilibrated in a mixture of 5% Heparin Buffer 1 (20 mM HEPES pH 7.5, 5% glycerol, 1 mM TCEP) and 95% Heparin Buffer 2 (20 mM HEPES pH 7.5, 1M NaCl, 5% glycerol, 1 mM TCEP). The column was then washed with 5 CV of this mixture of Heparin Buffer 1 and 2 followed by a gradient over 12 CV from 5% to 100% Heparin Buffer 2. Fractions containing polΔcat-prim were then pooled and concentrated using a 10 kDa MWCO spin filter, centrifuged, and applied to a S200 Increase column (Cytiva) equilibrated in SEC buffer (20 mM HEPES pH 7.5, 150 mM NaCl, 5 mM MgCl_2_, and 1 mM TCEP). Protein was eluted in 1.2 CV of this buffer, and fractions containing PolΔcat-prim were pooled and concentrated. Aliquots were then used immediately or flash frozen in liquid nitrogen and stored at - 80 °C. PolΔcat-primΔ2C was purified in the same manner while the polΔcat-primΔ1 construct lacking PRIM1 was purified from a single cell pellet.

### Sample Preparation and Crosslinking

5’-triphosphorylated RNA substrates were synthesized using as described previously.^46^ DNA oligonucleotides were purchased from Integrated DNA technologies and resuspended in MES buffer (10 mM MES pH 6.5, 40 mM NaCl). To generate RNA-primed templates, 66 uM RNA primer and 60 uM DNA templates were annealed in MES buffer by heating to 85 °C then slowly cooled to 4 °C. Protein substrate complexes for SAXS and EM experiments were prepared by incubating protein and primed substrate at a 1:1.1 ratio in binding buffer for 30 min. For crosslinking, bis(sulfosuccinimidyl)suberate (BS3) was purchased from Thermo fisher. A 10 mM stock solution was prepared immediately before use. Crosslinking reactions were performed using 2 mM BS3, 5 uM protein, and 5.5 uM substrate in SEC buffer, incubating at room temperature in the dark for 30 minutes. The reaction was quenched using 50 mM TRIS buffer. The reaction mixture was then subjected to Size-Exclusion Chromatography to remove higher order oligomeric species. Production of crosslinked products was monitored by SDS-PAGE.

### Microscale Thermophoresis

RNA/DNA binding reactions were conducted in the dark in binding buffer [20 mM HEPES•NaOH (pH 7.5), 150 mM NaCl, 5 mM MgCl2, 1 mM TCEP, 1mM AMPCPP and 0.05% (vol/vol) Tween-20] with 10 nM Cy5-labeled TriP 9mer primer-31mer template substrates. A 9mer RNA primer was used in lieu of 8mer to maximize retention of the primer on the template at the low concentration and near room temperature conditions in which the experiments were performed. Serial dilutions (1:1) of protein (2.5 μM to 0.61 nM) were prepared in binding buffer and combined with an equal volume of 20 nM Cy5-labeled primer-template. Samples were mixed and incubated on ice for 10 min, then at 17 °C for 10 min, and centrifuged at 20k ×g at 4 °C for 5 min prior to loading into standard-coated capillary tubes (NanoTemper, GmbH, Munich, Germany). Microscale thermophoresis data were collected on a Monolith NT.115 (NanoTemper) instrument. All experiments were conducted using the red fluorescence laser at 60% power (λ_excitation_: 600 – 640 nm; λ_emission_: 660 – 720 nm) and the infrared laser at medium power (λ = 1480 nm). Temperature-related intensity changes (TRIC) were performed at 17 °C with a pre-IR phase of 3 sec, followed by an IR-on phase of 20 sec, and a post-IR phase of 1 sec. Data quality was tracked by monitoring fluorescence quenching, photobleaching, protein aggregation, and adsorption for each experiment. Experiments were performed in triplicate. For each protein-DNA binding titration, the normalized fluorescence values were tracked as parts per thousand using the equation:

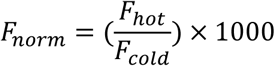

where F_hot_ corresponds to the mean fluorescence intensity from 0.5-1.5 seconds and F_cold_ to the mean fluorescence intensity from -1.0 – 0.0 seconds before irradiation and converted to ΔFnorm:

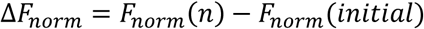

Dissociation constants were derived from fits to the data using a Law of Mass Action binding model:

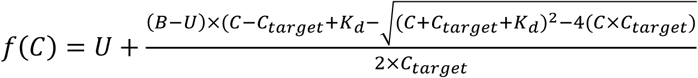

where f(C) is the fraction bound at a given protein concentration C, C_target_ is the concentration of fluorescently labeled oligonucleotide, B is the F_norm_ signal of the complex, U is the F_norm_ signal of the protein alone, and K_d_ is the dissociation constant.

### Small Angle X-ray Scattering

Scattering data were collected at the ALS beamline 12.3.1 LBNL Berkeley, California. The X-ray wavelength *λ* was 1.03 Å and the sample-to-detector distance was set to 1.5 m resulting in scattering vectors, *q*, ranging from 0.01 to 0.5 Å^−1^.^62^ The scattering vector is defined as *q* = 4π sinθ/*λ*, where 2θ is the scattering angle. For SEC-SAXS experiments, 60 μl of sample (25 uM) was loaded onto a Shodex 802.5 SEC column connected to a multi-angle light scattering system and the protein complex was eluted in SEC buffer.^63^ The eluent was split between SEC-MALS and SAXS channels. 3.0-s X-ray exposures were collected continuously during a ∼35 min elution. The SAXS frames recorded prior to the protein elution peak were used as buffer blanks to subtract from all other frames. The subtracted frames were examined by radius of gyration (Rg) and scattering intensity at q = 0 Å−1 (I(0)), derived using the Guinier approximation I(q) = I(0)*e*−*qRg*2/3 with the limits q*Rg <1.5. I(0) and Rg values were compared for each collected SAXS curve across the entire elution peak. The elution peak was mapped by plotting the scattering intensity at q = 0Å−1 (I(0)), relative to the recorded frame. Uniform Rg values across an elution peak represent a homogenous assembly. Data quality was assessed by: (i) fitting of the data in the Guinier region; (ii) good agreement between Rg calculated by the Guinier method and that based on P(r) distribution; (iii) molecular weight estimates derived from the data (Porod volumes) that were consistent with the expected size of the protein.^64^ The SAXS data were processed using Scatter 4.0 and PRIMUS.^65,66^ Back-calculated scattering data from coordinates extracted from structures available in the PDB were generated using FoXS.^52,67^ High-Throughput SAXS was performed using the same buffer and protein concentrations as SEC-SAXS.^68^ An array of ssDNA (0.2-2 mM) and GMPCPP (0.1-10 mM) concentrations were screened and monitored for major changes in the Mid-Q range of the Kratky plot that is highly sensitive to changes in shape. The concentration with the highest magnitude shift in the Mid-Q region without evidence of aggregation was further processed and compared to equivalent data collected for the free protein and a back-calculated scattering curve determined from a theoretical model of the configuration of PRIM2C that is primed for initiation (see Modeling section).

### EM sample preparation and data collection

Negative stain EM samples were prepared by diluting freshly purified polΔcat-prim to a final concentration of 50 nM in SEC buffer. 400 mesh carbon film copper grids (Electron Microscopy Sciences) were glow discharged for 2 minutes for non-crosslinked samples and 3 minutes for crosslinked samples. 2.5 uL of sample was applied to the grid and incubated for 1 minute, then blotted using filter paper, washed for 5 seconds in DI water, blotted, washed for 5 seconds in Uranyl formate stain, blotted, then stained for 90 seconds in Uranyl Formate solution. Grids were screened using a Morgagni 100 kV screening microscope. Micrographs were collected using a FEI 200 kV TF20 microscope.

### EM data processing

Initial data processing was performed in Sphire 1.9. Particle picking was performed with Cryolo yielding 80,000 – 100,000 particles per dataset.^69^ 2D classification was then performed in sphire using Iterative stable alignment clustering (ISAC).^70^ Particles were imported into cryosparc for further processing.^71^ Multi-class ab initio reconstruction and heterogeneous refinement was performed, and particles from classes displaying incomplete or irregular density were removed before submitting to further rounds of multi-class ab initio reconstruction and heterogeneous refinement (Figure S6).

### Modeling

Models of the tetramer core were generated by extracting coordinates from the PDB entries of eukaryotic pol-prim (5EXR, 8DOB, 7UY8, 7U5C, 8D9D, 8G9F) and rigid-body docked models from EMD entries (29888, -29889, -29891).^28,43,46,47,50^ Only residues from POLA2, POLA1C, PRIM2N, and PRIM1 were kept, and structures were aligned to POLA2/POLA1C for visualization. Two-state SAXS models were generated in Multi-FoXS.^52^

Models of polΔcat-prim were generated by extracting coordinates from the PDB entries of pol-prim in which PRIM2C was visible (PDB:5EXR, 7U5C, 8G9F, 8FOC, 8FOD, 8FOE, 8FOH, and 8D0K).^28,43,45–47^ Only residues from POLA2, POLA1C, PRIM2, and PRIM1 were kept and structures were aligned to PRIM1. For direct comparison to SAXS data, missing residues were added using MODELLER.^72,73^

BILBOMD was used to generate additional single and multiple configuration models of polΔcat-prim that are consistent with the experimental SAXS data.^74^ For these calculations, residues 245-265 and 455-509 in PRIM2 were defined as flexible.

Models of the RNA initiation and elongation configurations of polΔcat-prim were prepared by manually aligning PRIM1 from PDB:6R5D to PRIM1 to the tetramer core of PDB:5EXR.^54,75^ The structure of PRIM2C bound to substrate from PDB:8D9D was manually aligned such that the 3’ terminus of the RNA primer was aligned with the ATP molecule from PDB:6R5D for nucleophilic attack.^47^ All but 8 nucleotides for elongation or all but 1 nucleotide for initiation were retained from the structure of the RNA-primed template bound to PRIM2C. ^75^

As an aide to interpreting the SAXS data of polΔcat-prim in the presence of RNA initiation substrates, we performed a series of additional simulations to estimate to what extent a shift in population would be detectable using two models: one of polΔcat-prim in an ‘open’ configuration with PRIM2C away from PRIM1 and the other our model of the initiation configuration with PRIM2C aligned to PRIM1 (Figure S5D). These involved back-calculating the scattering profiles for different mixtures of the two configurations and comparing to the experimental data. From these simulations we concluded that a precision of ±20% could be obtained in estimating the ratio of one configuration relative to the other.

Models were initially aligned to EM density using the Fit in Map tool of ChimeraX to fit the teteramer core. Subsequently, atomic models were manually adjusted by treating the PRIM2C:PRIM1:substrate as an independent module and PRIM2N:POLA1C:POLA2 as another independent module on the basis of the large degree of conformational heterogeneity observed between the two in our SAXS and modeling studies.

## Acknowledgements

This work was funded by the National Institutes of Health (2T32GM008320-31 and R35GM118089 to WJC and R35GM136401 to BFE).

Small-Angle X-ray scattering was conducted at the Advanced Light Source (ALS), a national user facility operated by Lawrence Berkeley National Laboratory on behalf of the Department of Energy, Office of Basic Energy Sciences, through the Integrated Diffraction Analysis Technologies (IDAT) program, supported by DOE Office of Biological and Environmental Research. Additional support comes from the National Institute of Health project ALS-ENABLE (P30 GM124169) and a High-End Instrumentation Grant S10OD018483.

EM data were acquired at the Center for Structural Biology Cryo-EM Facility at Vanderbilt University. We acknowledge the use of the Glacios cryo-TEM, which was acquired by NIH grant S10 OD030292-01.

We thank Melanie Ohi and Jason Porta for their input in collecting and processing EM data.

## Author Contributions

JJC designed and performed experiments, analyzed results, and authored the manuscript. EAM contributed to experimental design, produced Tri-phosphorylated RNA primers, contributed to interpretation of results, and edited manuscript. LES developed cloning, expression, and purification protocols for proteins used. BFE contributed to interpretation of results and edited the manuscript. WJC oversaw the study, contributed to design and analysis of experiments, and wrote and edited the manuscript.

## Data Availability

Scattering data for polΔcat-prim, polΔcat-primΔ2C, polΔcat-prim in the presence of RNA Initiation factors, and polΔcat-prim in the presence of an RNA elongation substrate are deposited in the SASDB under accession codes DRAFT ID: 5126, 5127, 5128, and 5129 respectively. Negative stain EM data for crosslinked polΔcat-prim without substrate are deposited in the EMD under accession codes D_1000275627 for configuration 1 and D_1000275628 for configuration 2 while data of polΔcat-prim in the presence of an RNA elongation substrate are deposited under accession codes D_1000275599, D_1000275600, and D_1000275625 for configurations one, two, and three respectively.

---

**Table S1.**
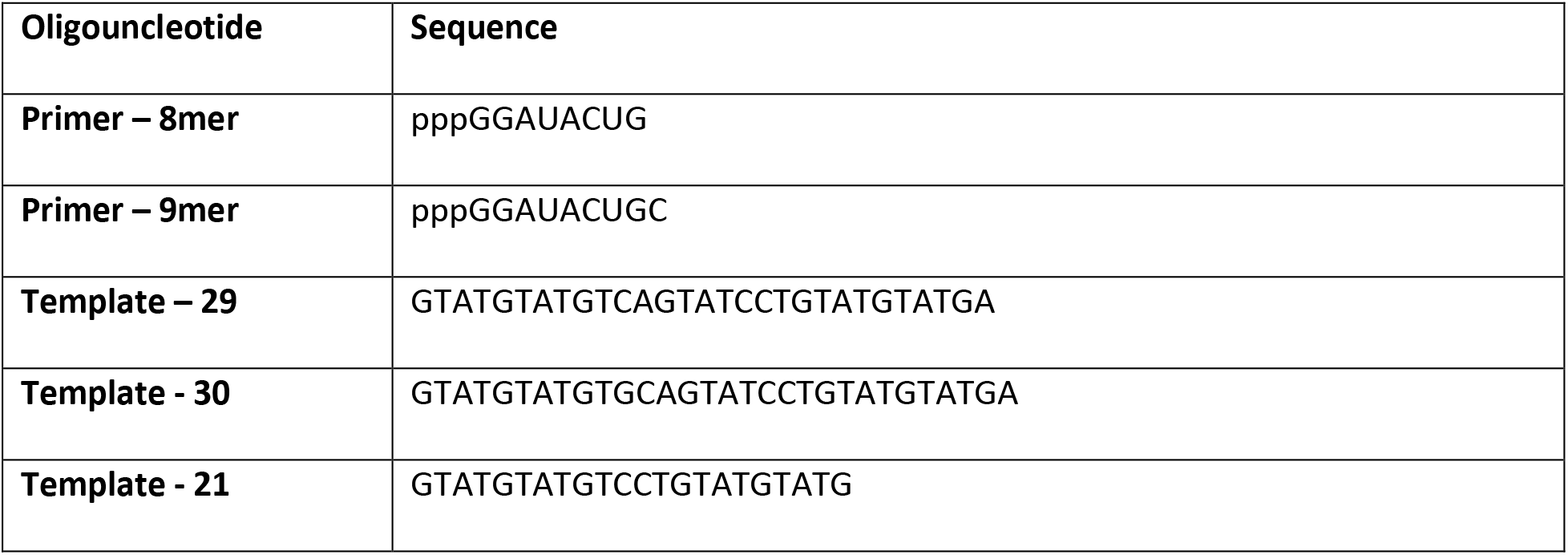
Oligonucleotides.

