## Supplemental Material for "Flexibility and distributive synthesis regulate RNA priming and handoff in human DNA polymerase α-primase"

**Supplementary Materials**

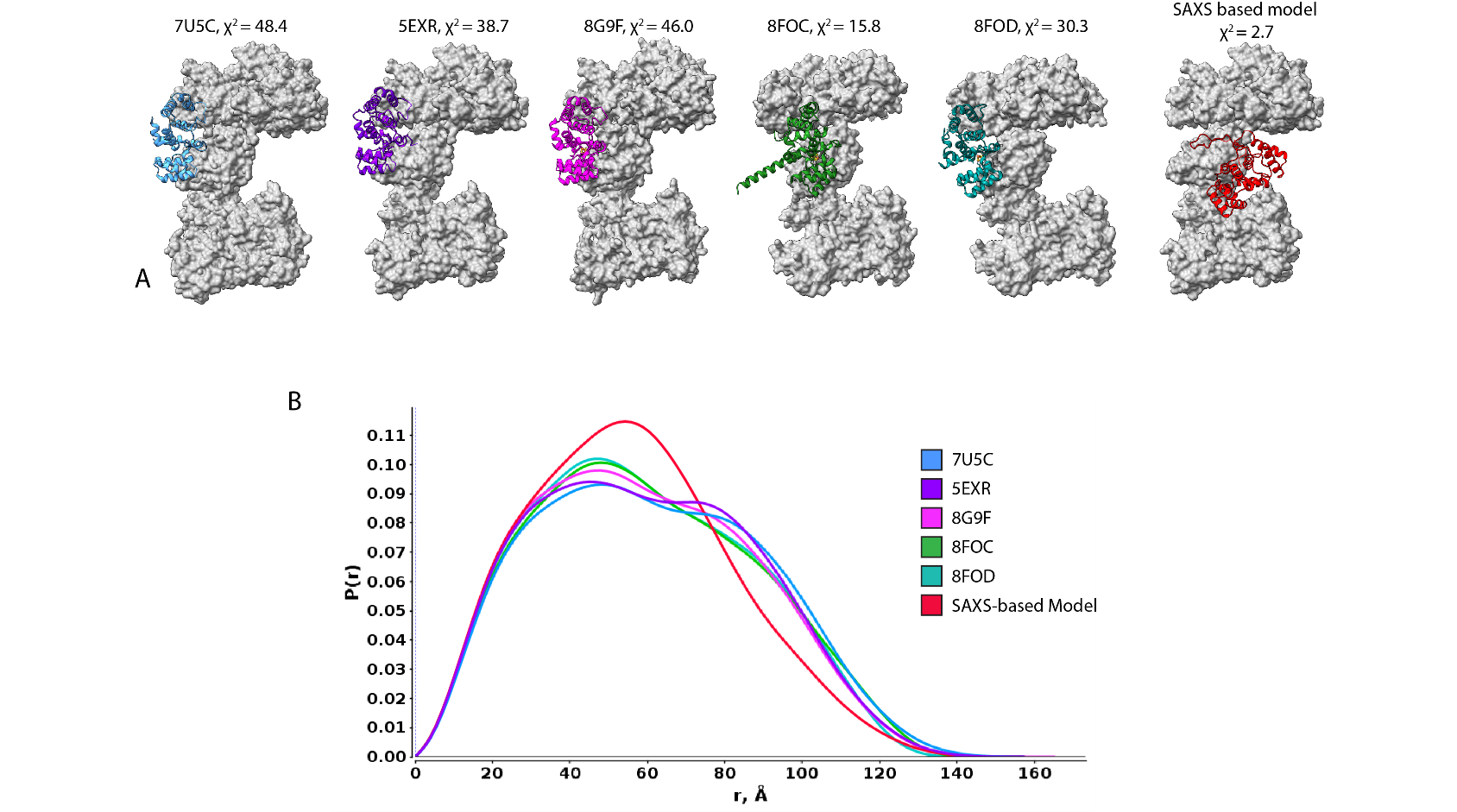

**Figure S1. Modeling Configurational Flexibility of PRIM2C in the absence of substrate.** A) Models of polΔcat-prim extracted from published structures of eukaryotic pol-prim in the absence of substrate and a SAXS-based limited MD simulation model. In models of published structures, only atoms corresponding to polΔcat-prim were kept. Structures in which CST was present are marked with an asterisk. Tetramer core is depicted in gray surface representation while PRIM2C is highlighted in colored ribbon. Models are aligned to PRIM1. χ^2^ values for each model are in comparison to polΔcat-prim scattering data. B) back calculated distance distributions from the models in A.

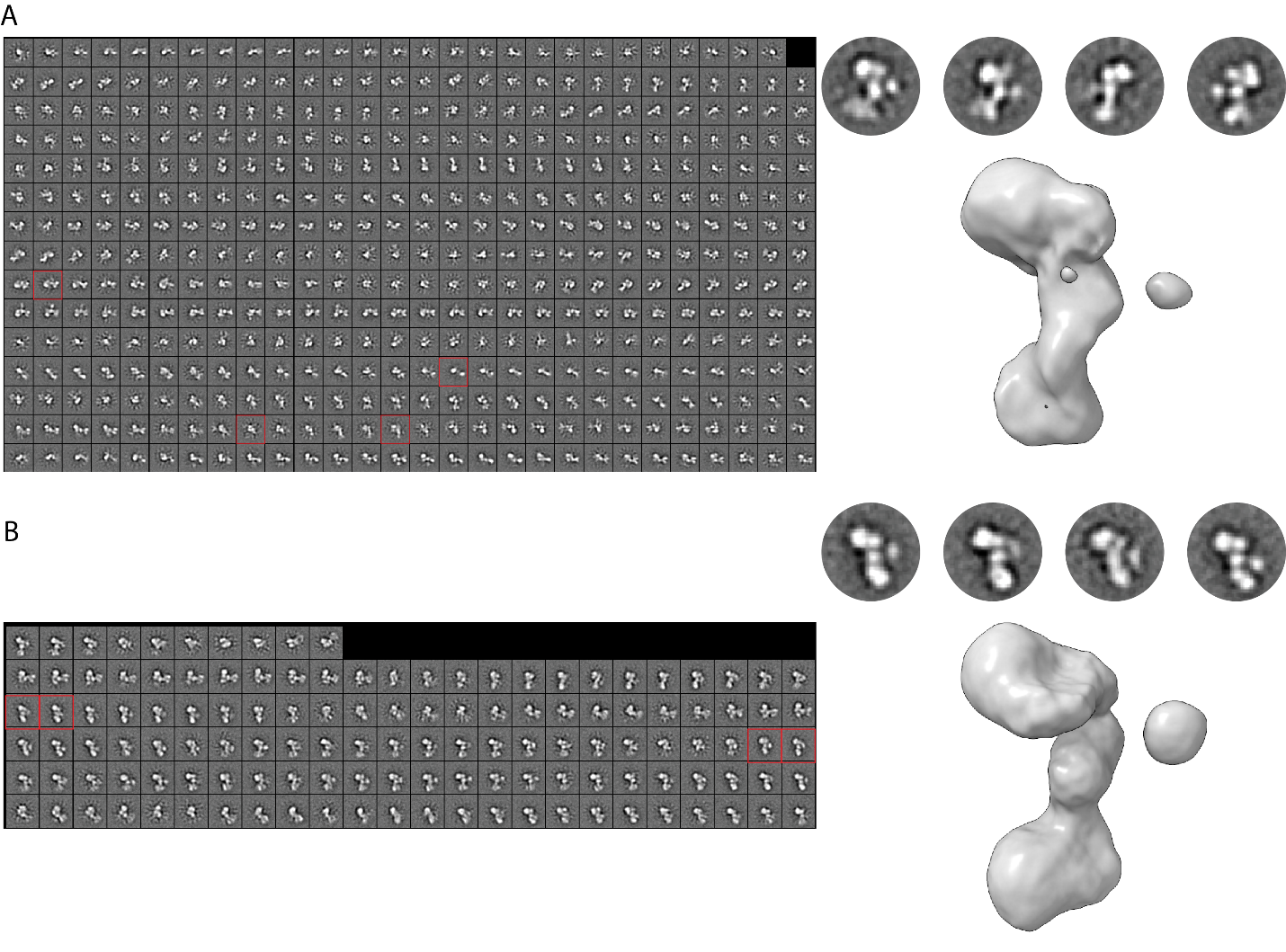

**Figure S2. Negative Stain EM without and with an RNA elongation substrate.** 2D class averages with selected 2D classes boxed in red and magnified at right, with consensus 3D reconstruction for polΔcat-prim (A) without and (B) with the RNA elongation substrate. 2D classes were created using ISAC in sphire while 3D reconstruction was performed in cryoSPARC.

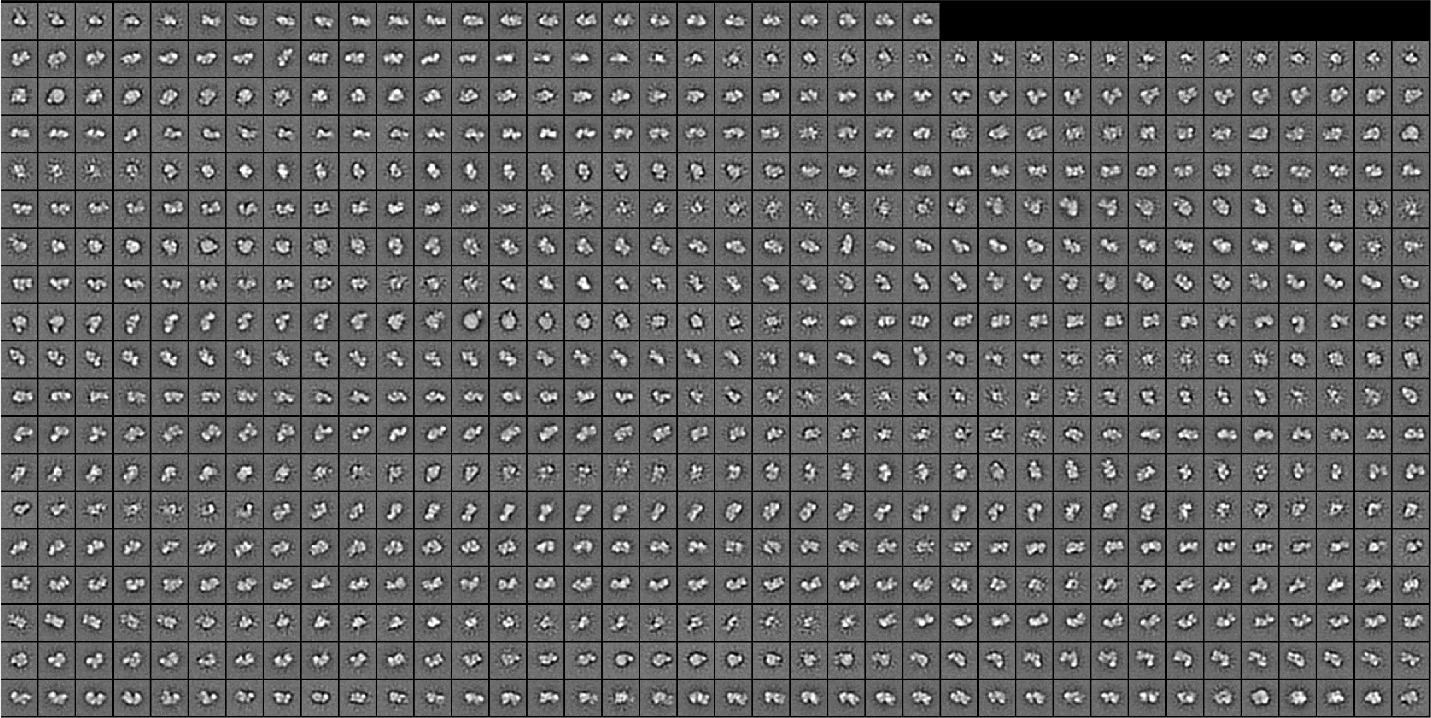

**Figure S3. 2D class averages of crosslinked no substrate polΔcat-prim.** 2D classes were created using ISAC in sphire. Red boxes indicate selected class averages depicted in Figure 4.

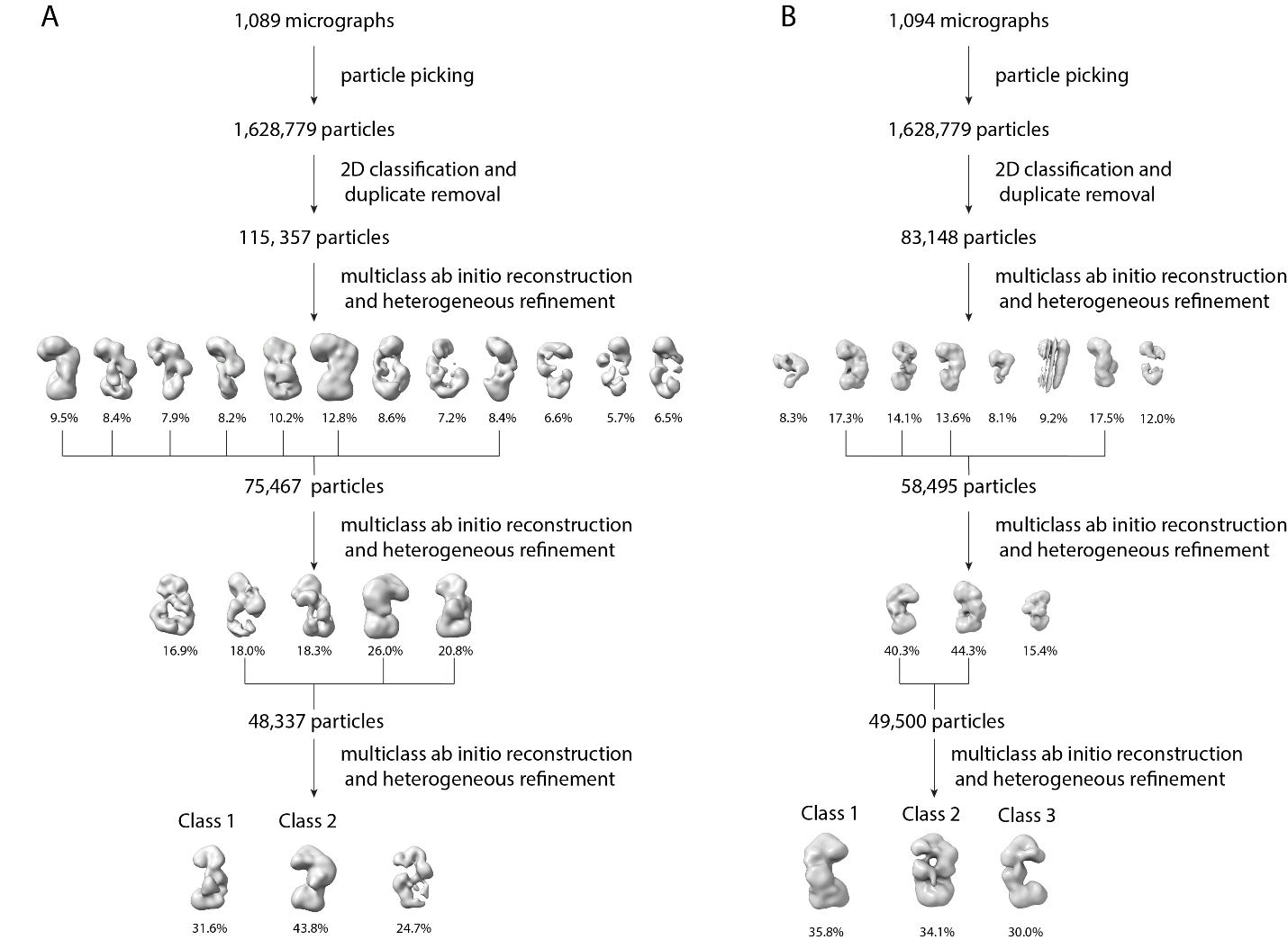

**Figure S4. Negative Stain EM data processing approach.** Initial particle picking was performed using crYOLO in sphire and box-particle picking in cryoSPARC. Subsequent steps were performed in cryoSPARC. A) No substrate, crosslinked. B) Elongation substrate, crosslinked.

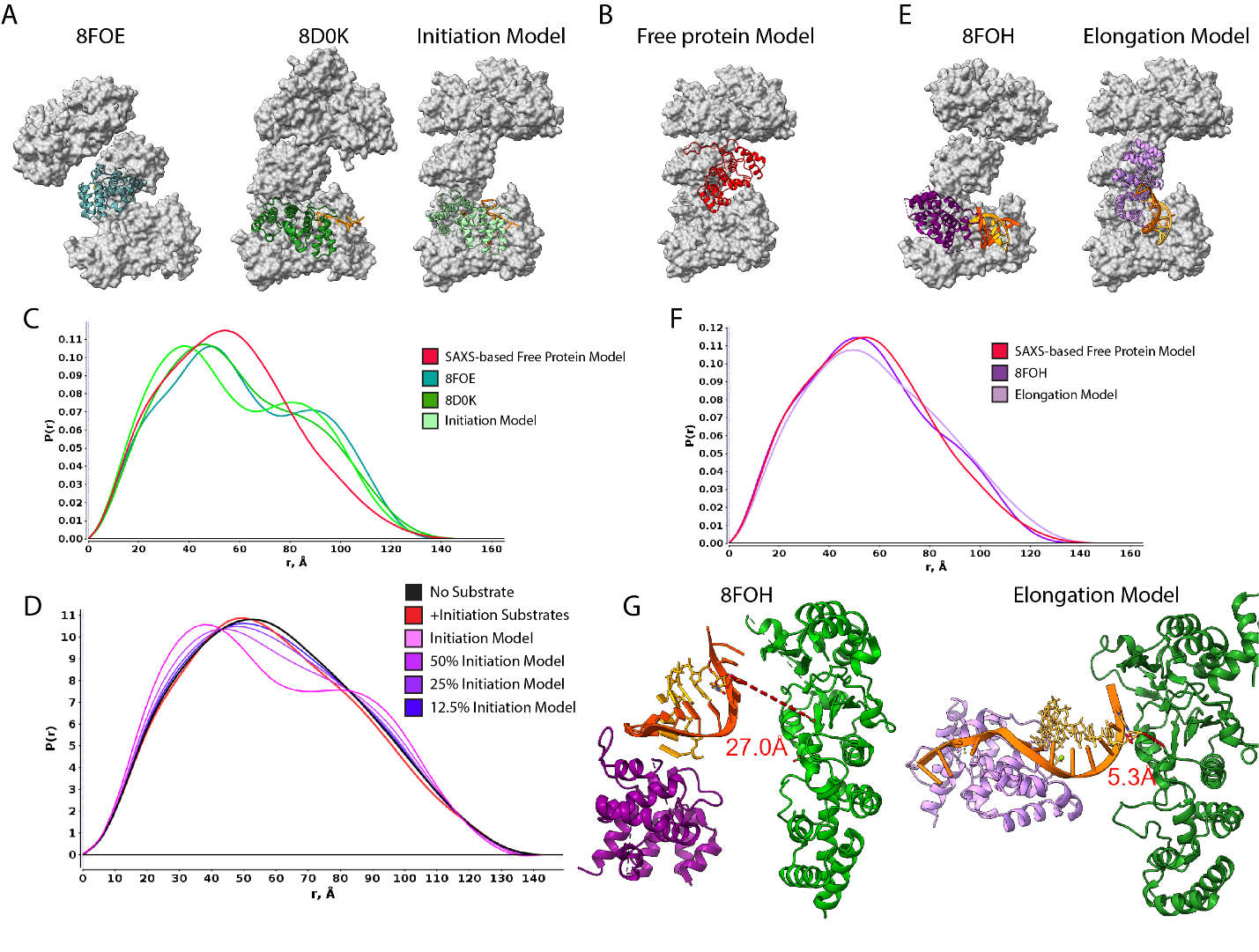

**Figure S5. Modeling configurational change of PRIM2C during RNA priming.** A) Models of polΔcat-prim in the presence of initiation substrate. Tetramer core is depicted in gray surface representation while PRIM2C is highlighted in colored ribbon. Models are aligned to PRIM1. B) No-substrate SAXS-based model of polΔcat-prim as shown in Figure S1 for comparison. C) back-calculated distance distribution from above atomic models. D) Comparison of atomic model of initiation complex (Fig S1, pink) with experimental data (red) and mixtures of free protein (black) and initiation model (purple to blue). E) Models of polΔcat-prim in the presence of RNA elongation substrate. Tetramer core is depicted in gray surface representation while PRIM2C is highlighted in colored ribbon. Models are aligned to PRIM1. F) back-calculated distance distribution from above atomic models of polΔcat-prim bound to RNA-primed template. G) Comparison of orientation of PRIM2C and PRIM1 in a recently published structure (PDB: 8FOH)^45^ and a manual model of RNA elongation. A large distance is observed between the catalytic ASP of PRIM1 and the 3’ Oxygen of the RNA primer in the published structure while in our manual PRIM1 engages the substrate for catalysis.

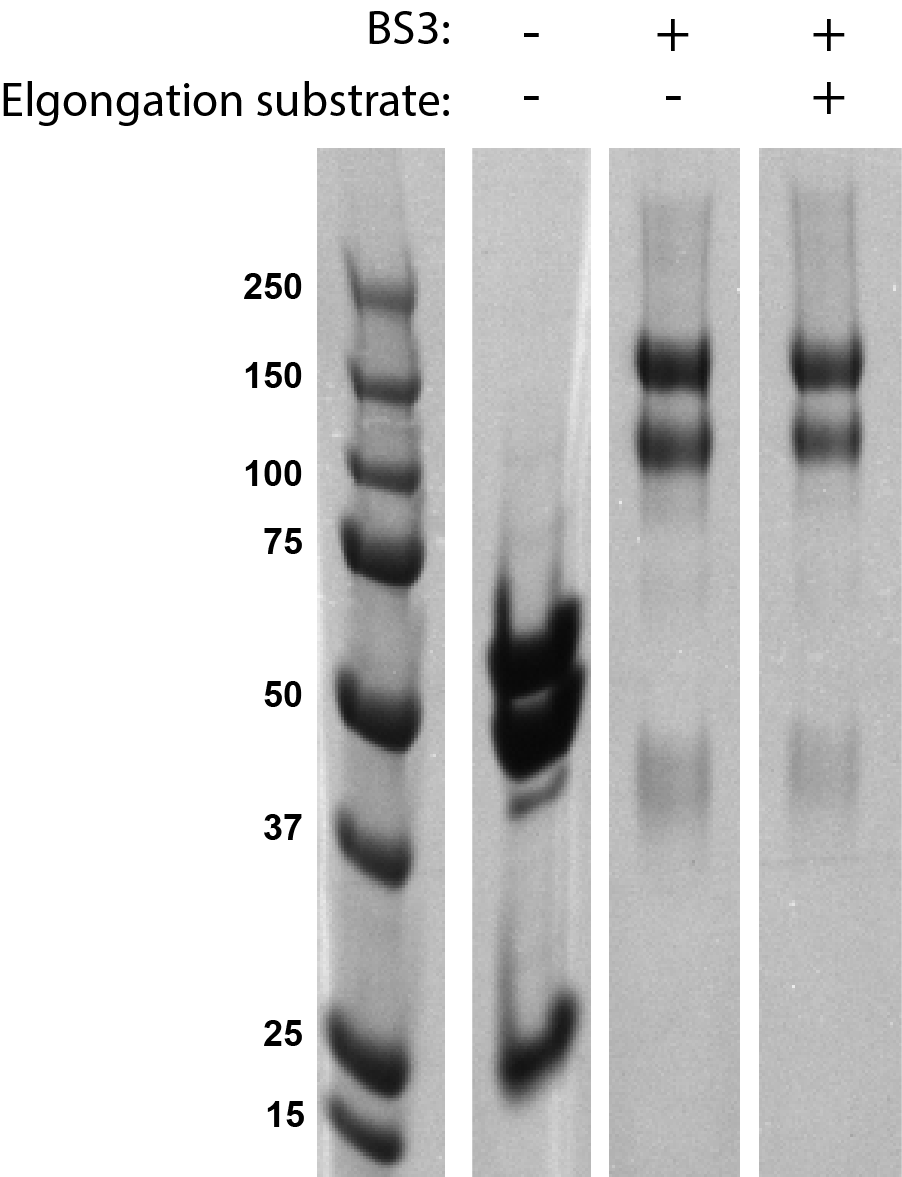

**Figure S6. BS3 crosslinking products analyzed by SDS-PAGE.** polΔcat-prim in the presence and absence of both BS3 crosslinker and RNA elongation substrate.

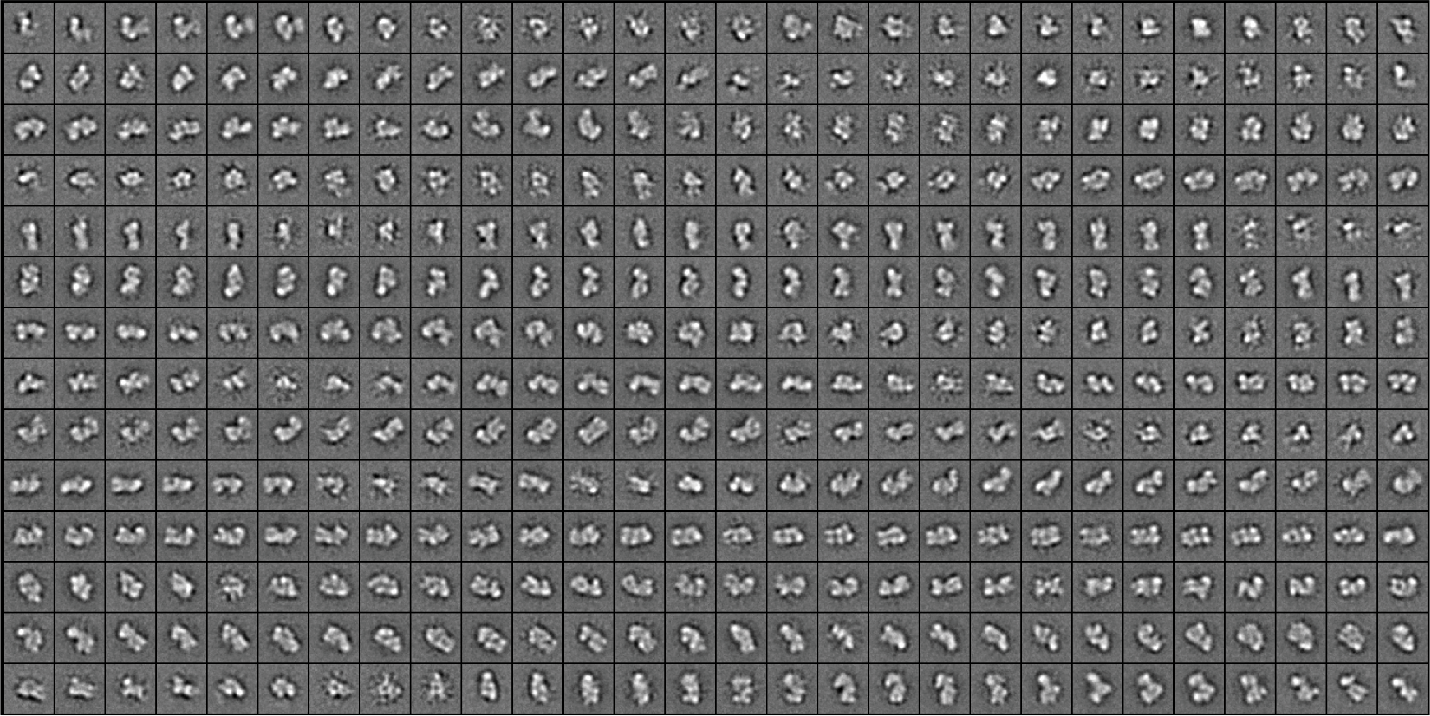

**Figure S7. 2D class averages of crosslinked polΔcat-prim in the presence of RNA elongation substrate.** 2D classes were created using ISAC in sphire. Red boxes indicate selected class averages depicted in Figure 4.

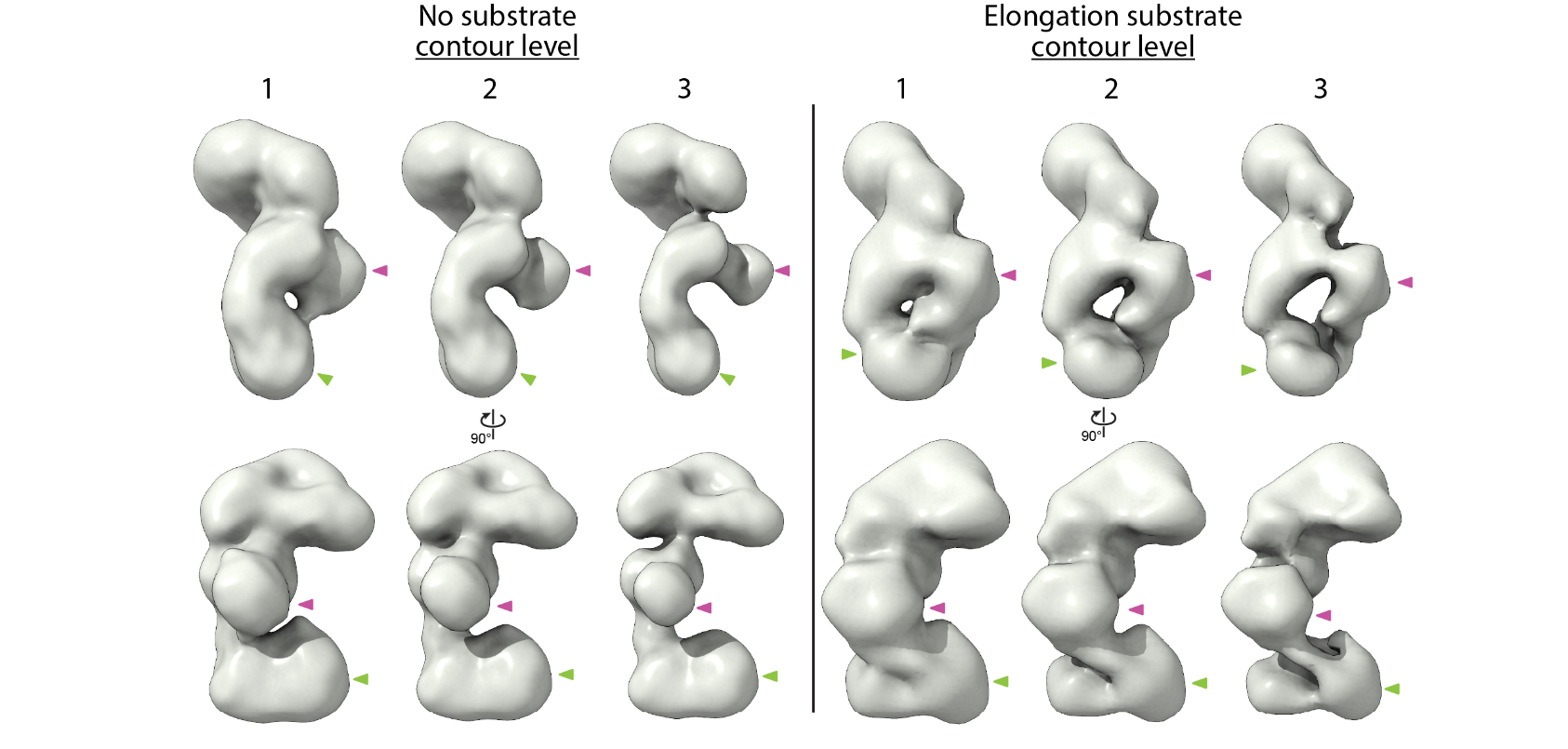

**Figure S8. Density Connects PRIM2C and PRIM1 after addition of RNA elongation substrates.** 3D reconstruction at 3 contour levels for A) class 1 no substrate and B) class 1 elongation substrate with PRIM1 (green) and PRIM2C (purple) indicated with arrows.
